# Targeting folate-dependent purine synthesis sensitizes melanoma cells to immune attack through suppressing glycolysis

**DOI:** 10.64898/2026.07.06.736685

**Authors:** Dimin Li, Miaomiao Hou, Sihong Wang, Xinyuan Wan, Haiqiong Wang, Yijie Han, Xi Liu, Chaojiang Cheng, Jian Zhang, Xiao Hu

## Abstract

Cytotoxic T lymphocytes (CTLs) play a central role in antitumor immunity; however, metabolic reprogramming within the tumor microenvironment often compromises their effector function, making metabolic targeting crucial for the improvement of T cell function. Folate-dependent purine synthesis, a core pathway sustaining the nucleotide pool, is highly activated in tumors, yet its role in regulating tumor immune sensitivity remains unclear. Here, by establishing a co-culture system of melanoma cells and human T Cell Receptor (TCR)-engineered T cells, we systematically evaluated the effects of folate-dependent purine synthesis inhibitors on tumor cell response to CD8+ T cell cytotoxicity. We found that inhibition of key enzymes such as methylenetetrahydrofolate dehydrogenase 2 (MTHFD2) and glycinamide ribonucleotide transformylase (GART) markedly enhanced tumor cell sensitivity to T cell killing, an effect also observed with exogenous nucleoside supplementation. Mechanistically, inhibition of folate-dependent purine synthesis suppresses glycolysis by downregulating critical glycolytic enzymes, thereby reducing lactate production. Reduction in lactate further weakens lactylation and stability of the immune checkpoint protein PD-L1. In parallel, impaired purine synthesis disrupts uridine metabolism, blocks ribose salvage, and distally influences glycolysis. Collectively, our study identified the folate-dependent purine synthesis – glycolysis axis as key regulator of tumor immune response and highlights metabolic targeting as a promising strategy to improve cancer immunotherapy.

## Introduction

Cytotoxic T lymphocytes (CTLs), particularly CD8+ T cells, specifically recognize antigen presenting cells (APCs) through the interaction between T cell receptors (TCRs) on T cells and antigenic peptides presented by class I major histocompatibility complex (MHC-I) on APCs. Recognition activates T cells to perform effector function by secreting pro-inflammatory cytokines (such as TNF-α and IFN-γ) and releasing cytotoxic molecules (such as perforin) to directly eliminate tumor cells, which are typical APCs.^1^ In recent years, tumor metabolism has emerged as a critical factor regulating CTL-mediated tumor immune response by modulating both tumor intrinsic properties and tumor microenvironment (TME).^2^ Among these metabolic pathways, folate-dependent purine synthesis, which integrates one-carbon (1C) metabolism and de novo purine synthesis, is essential for tumor growth but its involvement in tumor immunity is relatively underexplored.

Functioning as a core metabolic network, 1C metabolism integrates multiple nutrients to support nucleotide biosynthesis, amino acid metabolism, redox homeostasis, and epigenetic regulation.^3,4^ Elevated 1C metabolism is a hallmark of metabolic reprogramming in many cancers. Traditional chemotherapeutic agents, such as methotrexate (MTX) and 5-fluorouracil (5-FU), have been used to target this pathway in cancer treatment for nearly 70 years.^5^ More recently, 1C metabolism inhibitors have been found to exert immunomodulatory effects through regulating immune cell metabolism. For instance, inhibitors of serine hydroxy-methyltransferase (SHMT) limit effector T cell expansion by restricting serine metabolism, thereby impairing immune responses while inhibitors of S-adenosylhomocysteine hydrolase (ACHY) or adenosine deaminase (ADA) suppress CD4+ T cell activation via the adenosine metabolic pathway.^6,7^ Nevertheless, evidence showing how modulating 1C metabolism in tumor cells alters their immune response is still lacking.

Importantly, 1C metabolism and de novo purine synthesis is tightly interconnected with glycolysis. The glycolytic intermediate 3-phosphoglycerate feeds into the serine synthesis pathway to provide carbon for 1C metabolism.^8-10^ Glucose-6-phosphate from glycolysis enters the oxidative pentose phosphate pathway (oxPPP) to generate phosphoribosyl diphosphate (PRPP), together with 10-formyl-tetrahydrofolate (10-f-THF) from 1C metabolism, to support de novo purine synthesis.^11,12^ Reciprocally, nucleotides support glycolysis through ribose salvage involving non-oxidative pentose phosphate pathway (non-oxPPP).^11,13,14^ Given such connections, chemotherapeutic agents aimed at targeting 1C metabolism or de novo purine synthesis should interfere with glycolysis, whose increase not only supplies energy and biosynthetic precursors but also causes lactate accumulation and microenvironment acidification – a phenomenon known as the Warburg effect.^15^ Therefore, tumor glycolysis is a significant contributor to the immune suppressive TME where CD8+ T cells are often found functionally impaired,^16,17^ and targeting tumor glycolysis should sensitize tumor cells to CD8+ T cell killing. However, whether targeting 1C metabolism or de novo purine synthesis in tumor cells benefit T cell immune attack similarly is unknown. In this study, we employed a co-culture system of melanoma cells and CD8+ T cells to systematically evaluate the effects of 1C metabolism and nucleotide synthesis inhibitors on tumor cell sensitivity to T cell cytotoxicity, with the underlying metabolic mechanisms explored. Our findings reveal that inhibition of 1C metabolism by a methylenetetrahydrofolate dehydrogenase 2 (MTHFD2) inhibitor or de novo purine synthesis by a glycinamide ribonucleotide transformylase (GART) inhibitor enhances tumor cell sensitivity to T cell-mediated killing by suppressing glycolysis both transcriptionally and metabolically. These results highlight folate-dependent purine synthesis as a promising metabolic target for optimizing melanoma immunotherapy.

## Results

### MTHFD2 inhibition promotes tumor cell susceptibility to CD8+ T cell killing

We previously established a tumor cell killing model using HLA-A*02:01-restricted TCRs 1G4 and its activity-enhanced variant 1G4 α95LY that specifically recognize the tumor antigen peptide SLLMWITQC from the protein New York esophageal squamous cell carcinoma 1 (NY-ESO-1_157–165_) encoded by the gene *CTAG1A/B*.^18^ We aimed at identifying drugs that could potentially sensitize tumor cells to cytotoxic CD8+ T cell killing. The melanoma cell line A375 expresses both the cognate MHC class I molecule and low levels of NY-ESO-1, and thus it serves a perfect cell line to monitor the cytotoxic activity of T cells expressing the corresponding TCRs. A375 cells were pretreated with individual drugs from a targeted diversity library for 24 hours before they were confronted with T cells. Among the 2,376 drugs tested, approximately 19 drugs came out as the most effective ones in stimulating T cells, with an elevation of IFN-γ secretion for more than 1.5-fold in each of the two screening assays (Figure 1A). Most of these drugs that show effectiveness target components in cell cycle control, epigenetic regulation, and TGFβ signaling (Figure 1B). As a validation of our approach, the screening also revealed MDM2 inhibitor and HDAC inhibitors, which have been reported to facilitate cancer immunotherapy.^19,20^ The drug DS18561882, a selective inhibitor of MTHFD2, came to our attention since MTHFD2 is a critical mitochondrial enzyme involved in tetrahydrofolate cycle, also known as 1C metabolism. Due to its importance in de novo purine biosynthesis, which is often overactive in tumor cells, 1C metabolism has long been a target for cancer chemotherapy.^21^ We were particularly interested in exploring how its inhibition is connected to the enhancement of TCR-T cell-mediated tumor cell killing, as this holds promise for developing novel strategies using 1C metabolism drugs for the improvement of adoptive T cell transfer therapy.

**Figure 1.**
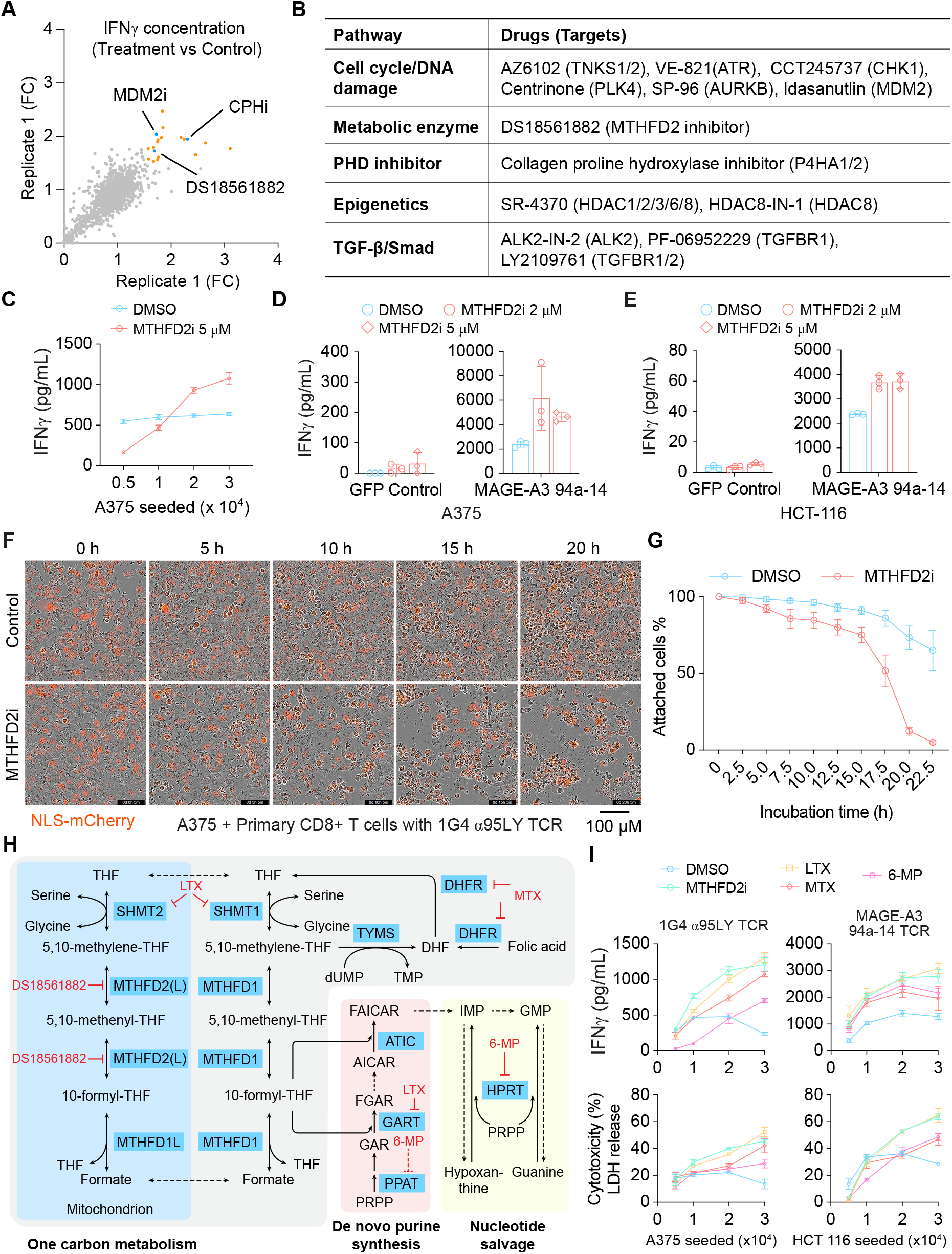
MTHFD2 inhibition promotes CD8+ T cell activation and tumor cell killing. (A) Drug screening results showing fold changes in IFN-γ secretion (treatment versus control) in two independent screening replicates. Each dot represents one compound. Selected hits are indicated. (B) Classification of the most effective inhibitors identified in this study and their corresponding targets and signaling pathways. (C) IFN-γ secretion in the co-culture supernatants. A375 cells were seeded at indicated densities to grow for 24 h, then treated with DMSO or MTHFD2i (5 μM) for another 24 h, followed by co-culture with primary CD8+ T cells expressing 1G4 α95LY TCR for 24 h. (D, E) IFN-γ levels in the supernatants after co-culture of MTHFD2i-pretreated A375 (D) or HCT 116 (E) cells with human primary CD8+ T cells transduced with control GFP or MAGE-A3 94a-14 TCR, as indicated. MTHFD2i was used at 2 μM or 5 μM. (F) Live-cell imaging showing the dynamics of TCR-T cell-mediated killing of A375 cells pretreated with DMSO or MTHFD2i (5 μM) for 24 h. (G) Percentage of attached A375 cells over time during co-culture. A375 cells were treated with DMSO or MTHFD2i (5 μM) for 24 h before T cells were added. (H) Schematic illustration of 1C metabolism, de novo purine synthesis, and nucleotide salvage pathways. Inhibitors targeting DHFR (methotrexate, MTX), MTHFD2 (DS18561882), GART and SHMT1/2 (lometrexol, LTX), HPRT or indirectly PPAT (6-mercaptopurine, 6-MP) are highlighted. (I) IFN-γ secretion (upper panels) and LDH release (lower panels) in the co-culture supernatants. A375 or HCT 116 cells were seeded at indicated densities to grow for 24 h, then treated with DMSO, MTHFD2i (5 μM), LTX (2 μM), MTX (2 μM), or 6-MP (100 μM) for another 24 h, followed by co-culture with human primary CD8+ T cells expressing 1G4 α95LY TCR or MAGE-A3 94a-14 TCR for 24 h. Elisa data are representative of 3 independent experiments unless otherwise indicated. Data are presented as mean ± SD.

We first validated our findings by performing co-culture, where A375 cells were treated with DS18561882 (hereafter referred to as MTHFD2i) 24 hours prior to the addition of human primary CD8+ T cells expressing exogenous 1G4 α95LY TCR. Consistent with the screening result, MTHFD2i pretreatment led to a tumor cell density-dependent increase in IFN-γ secretion by CD8+ T cells, whereas this is not observed with DMSO pretreatment (Figure 1C). To examine whether the promotive activity of MTHFD2i pretreatment applies to other TCRs, we used HLA-A*01:01-restricted TCR 94a-14 that specifically recognizes the tumor antigen peptide EVDPIGHLY from Melanoma antigen family A, 3 (MAGE-A3).^22^ Both A375 and the human colon cancer cell line HCT 116 are HLA genotype matched and express MAGE-A3. Again, for both cell lines, different concentrations of MTHFD2 inhibition induced dramatic increase in IFN-γ secretion by 94a-14-expressing CD8+ T cells (Figures 1D and 1E), indicating the effect of MTHFD2 inhibition is conserved across different cancer cell types. Moreover, the enhancement is TCR-specific since control CD8+ T cells were not promoted for IFN-γ secretion.

To confirm that the enhanced stimulation of IFN-γ secretion is a faithful indication of CD8+ T cell cytotoxicity, we performed live-cell imaging to directly monitor the killing dynamics of T cells toward tumor cells. We included a nuclear localization signal (NLS)-tagged mCherry fluorescent protein to better visualize the integrity of tumor cells. Under the microscope, MTHFD2i-treated cells were more vulnerable to T cell killing, as revealed by the earlier appearance of lysed cells (Figure 1F, 5 h). Massive elimination was observed for drug-pretreated cells during the period of 10 h – 15 h post T cell adding, while cell death events were still limited in the control. By 20 h, almost all MTHFD2i-treated cells were lysed whereas more than 50% of DMSO-pretreated cells were still intact as indicated by nuclear mCherry signal (Figures 1F and 1G). These data demonstrate that MTHFD2 inhibition significantly potentiates T cell-mediated killing. To exclude potential off-target effects from MTHFD2i and establish a causal role of MTHFD2 inhibition in promoting tumor cell sensitivity, we generated two knockdown cell lines expressing distinct shRNAs targeting MTHFD2 (shMTHFD2-1/2) in A375 cells. Immunoblotting confirmed substantial reduction of MTHFD2 protein levels (Figure S1A). In the co-culture system, both knockdown cell lines phenocopied pharmacological inhibition, exhibiting target cell density-dependent enhancement of IFN-γ secretion by CD8+ T cells and lactate dehydrogenase (LDH) release (an indication of tumor cell lysis) by tumor cells (Figure S1B). Together, these data support the conclusion that suppressing MTHFD2 increases tumor cell susceptibility to antigen-specific CD8+ T cell killing.

As a mitochondrial enzyme in the tetrahydrofolate cycle, MTHFD2 catalyzes the two-step reversible conversion of 5,10-methylene-tetrahydrofolate (5,10-CH2-THF) into 10-formyl-tetrahydrofolate (10-f-THF), with the latter being a carbon donor facilitating de novo purine synthesis, which is a series of enzymatic reactions initiating from PRPP to form inosine 5’-monophosphate (IMP) (Figure 1H).^23,24^ To examine if the promotional effect is related to the block of MTHFD2-involved metabolic pathways, we included inhibitors targeting other members in 1C metabolism and de novo purine synthesis pathway (hereafter referred to as folate-dependent purine synthesis pathway), such as MTX, which interferes with dihydrofolate reductase (DHFR) in 1C metabolism, and lometrexol (LTX), which exhibits inhibitory effect on both GART in the de novo purine synthesis pathway and SHMT1/2 in the tetrahydrofolate cycle of 1C metabolism (Figure 1H). We also included 6-mercaptopurine (6-MP), a purine analogue that competes with the substrates of HPRT1, thereby inhibiting the nucleotide salvage pathway.^25^ It is notable that 6-MP can be converted into 6-methylmercaptopurine (6-meMP) by thiopurine methyltransferase (TPMT) or methylthioinosine monophosphate (meTIMP) by HPRT1 and TPMT. Both 6-meMP and meTIMP are inhibitors of PRPP amidotransferase (PPAT), thereby 6-MP indirectly inhibits de novo purine synthesis.^26,27^ Strikingly, tumor cells pretreated with these inhibitors also enhanced CD8+ T cell-mediated IFN-γ secretion, especially at high densities for both A375 and HCT 116 cells (Figure 1I), indicating folate-dependent purine synthesis inhibition might underlie the promotional effect. Importantly, these drugs display differential potency in sensitizing tumor cells, with MTHFD2i and LTX the strongest, followed by MTX and then 6-MP (Figure 1I). However, drug enhancement is hardly observed for the renal cell carcinoma cell line A498, with only 6-MP showing a mild effect on tumor cell lysis after co-culture with T cells (Figure S2). We reasoned that unlike A375 and HCT 116 cells where de novo purine synthesis plays a stronger role for tumor growth,^28^ cells showing high activities of nucleotide salvage, such as renal cell carcinoma cells,^29^ might be inefficient to sensitize tumor immune response after being pretreated with folate-dependent purine synthesis drugs.

### Drug-induced cell cycle inhibition does not account for enhanced tumor cell susceptibility to CD8+ T cell killing

Our findings that MTHFD2i and LTX can sensitize tumor cells for CD8+ T cell killing indicate these drugs are potentially suitable to be combined with adoptive T cell transfer therapy for cancer treatment. Thus, exploring the underlying mechanisms is of great significance. We found that drug-induced enhancement is not due to increased expression of HLA molecules or downregulated expression of inhibitory factors, such as *CD274* (encoding PD-L1), *CTLA-4*, and *LAG-3*, in tumor cells at the transcript level (Figure S3A). MTHFD2i treatment barely changed the surface expression of HLA-A, B, C based on fluorescence activated cell sorting (FACS) analysis (Figure S3B). We then performed detailed analysis of the transcriptome generated from A375 cells treated with MTHFD2i or LTX. Gene set enrichment analysis (GSEA) using cancer hallmark gene sets as reference revealed that MTHFD2 inhibition led to dramatic downregulation of cell cycle-related genes involved in or grouped as G2M checkpoint, E2F targets, and mitotic spindle assembly. In addition, cellular metabolism was notably altered as represented by compromised mTORC1 signaling and glycolysis. In contrast, the p53 pathway was significantly upregulated (Figure 2A). Similar results were obtained in the case of LTX treatment, with an even stronger suppression of glycolysis (Figure 2A). These data indicate that 1C metabolism inhibition by MTHFD2i not only causes cell cycle arrest but may also induce G1 checkpoint activation through upregulating p53 signaling, thereby extending the window for cell cycle progression preparation. At the same time, the unexpected suppression of glycolysis highlights that 1C metabolism inhibition directly impairs tumor energy supply, undermining the metabolic demands for rapid proliferation. More importantly, most of these alterations are shared between MTHFD2i and LTX, offering a conclusion that drug effects converge on de novo purine synthesis whose suppression results in shared downstream alterations.

**Figure 2.**
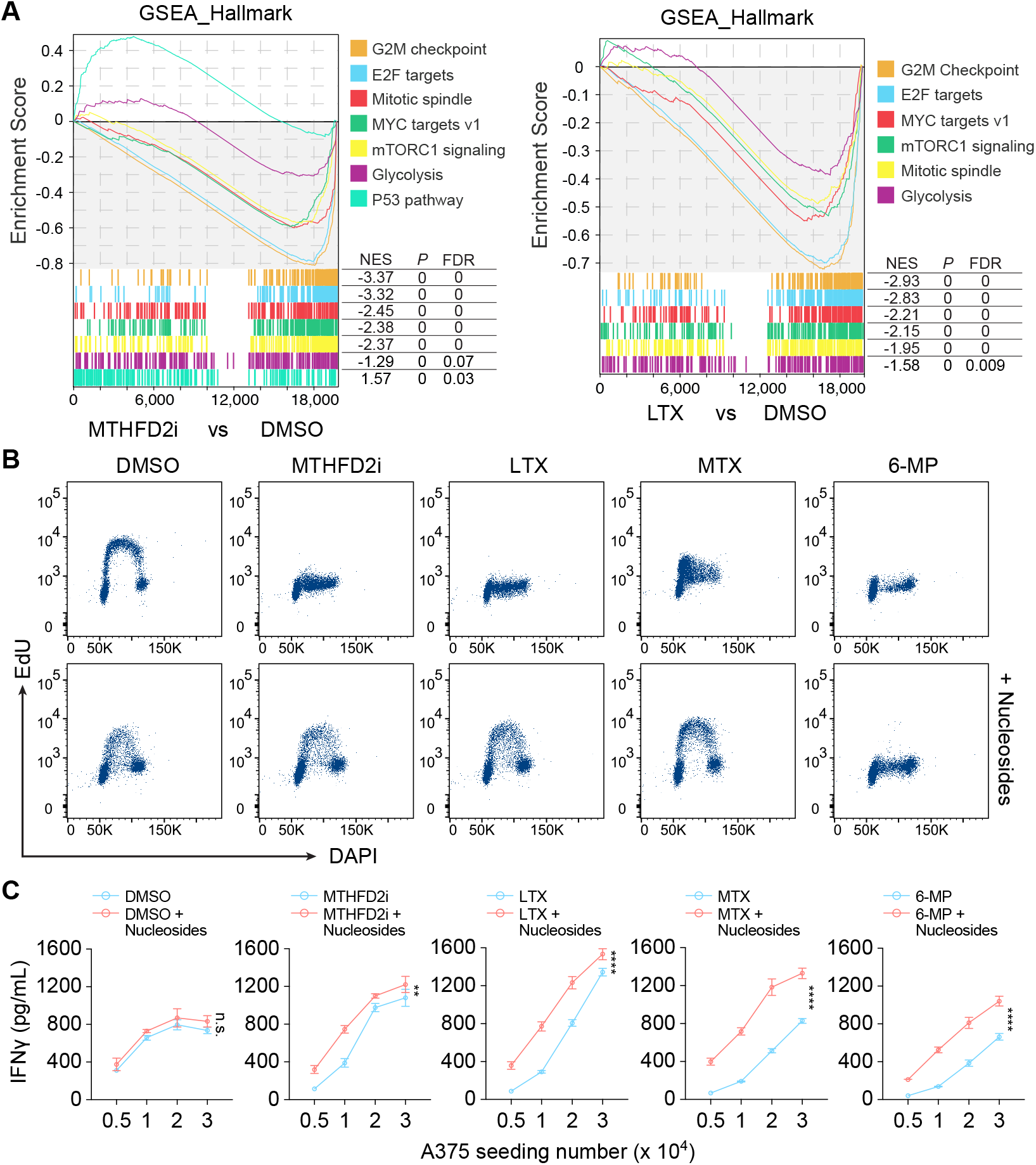
Enhancement of tumor immune response by MTHFD2i or LTX is maintained with nucleoside supplementation. (A) Gene Set Enrichment Analysis (GSEA) showing upregulated and downregulated pathways in A375 cells treated with MTHFD2i (5 µM) or LTX (2 µM) for 24 h. (B) Cell proliferation assessed by EdU incorporation and flow cytometry after treatment with DMSO, MTHFD2i (5 µM), LTX (2 µM), MTX (2 µM), or 6-MP (100 µM), with or without nucleoside supplementation. Nucleosides were added in a final concentration of 30 μM for cytidine, guanosine, adenosine, and uridine each, and 10 μM for thymidine. (C) IFN-γ secretion in the co-culture supernatants. A375 cells seeded at indicated densities were treated with DMSO, MTHFD2i (5 μM), LTX (2 μM), MTX (2 μM), or 6-MP (100 μM) supplemented with nucleosides for 24 h, followed by co-culture with 1G4 α95LY TCR-T cells. Data are representative of 3 independent experiments and shown as mean ± SD for each datapoint. Statistical analysis was performed using two-way ANOVA with corrections for multiple comparisons. n.s.: non-significant; **: *P* < 0.01; ****: *P* < 0.0001.

To determine whether cell cycle arrest in tumor cells is causative for the observed T cell response, we performed nucleoside supplementation driven by our hypothesis that de novo purine synthesis inhibition-induced deficiency in nucleotides is the reason DNA replication and cell cycle suspend. Indeed, the Fucci(SA)2.1 dual-reporter system,^30^ with AmCyan-hGem dictating S/G2/M phase and mCherry-hCdt1 dictating G1 phase, displayed great loss of AmCyan-positive cells, indicating block of the cells at G1/S phase transition and depletion of the cells in S phase (Figure S4A). Consistently, MTHFD2i, LTX, and MTX treated cells completely lost EdU incorporation, a sign of suppressed DNA synthesis, which can be relieved upon supplementation of mixed nucleosides (adenosine, guanosine, cytidine, thymidine, and uridine) (Figure 2B) but not adenine alone (Figure S4B). In line with failed rescue of DNA synthesis, adenine supplementation did not change the response of CD8+ T cells to drug-pretreated tumor cells as assessed by IFN-γ secretion during the co-culture (Figure S4C). Unexpectedly, although nucleoside supplementation managed to make DNA replication and cell proliferation resume, it boosted IFN-γ secretion (Figure 2C), making it impossible for cell cycle arrest as a causal factor in enhancing tumor immune sensitivity and implying that drug-induced persistent defects after nucleoside rescue might be responsible. Collectively, these findings indicate that mechanisms other than cell cycle arrest induced by MTHFD2i or LTX sensitizes tumor cells to CD8+ T cell killing.

### Folate-dependent purine synthesis inhibition induces metabolic stress and suppresses glycolysis

Having excluded cell cycle arrest as a major player, we turned to metabolic changes that are also pronouncedly impacted by these drugs according to our GSEA data. De novo purine synthesis inhibition suppresses the activity of mechanistic target of rapamycin (mTOR), a serine/threonine kinase in the mTOR complex 1 (mTORC1).^31,32^ We found that mTORC1 signaling was revealed as negatively influenced after MTHFD2i or LTX treatment in 375 cells (Figure 2A). Thus, we investigated the possibility of linking tumor mTORC1 signaling with T cell immunity.

We observed an enrichment of cytoplasmic ribosome-related gene expression after MTHFD2 inhibition (Figure S5A), implying a compensatory mechanism is activated to meet the demand for protein synthesis due to drug-induced mTOR activity inhibition. mTORC1 directly phosphorylates ribosomal protein S6 kinase (S6K) and eIF4E-binding protein (4EBP) to promote translation.^33^ Upstream, mTORC1 is positively regulated by a small GTPase Ras homolog enriched in brain (RHEB).^34,35^ Our data revealed that these inhibitors consistently led to marked decrease in phosphorylated S6K (p-p70-S6K) and RHEB protein, confirming that the activity of mTORC1 is strongly inhibited when either de novo or salvage nucleotide synthesis is blocked (Figure S5B). Supplementing exogenous adenine or nucleosides effectively rescued the reduction in p-p70-S6K and RHEB levels (Figure S5B), which agrees with previous reports that mTOR activity is regulated by the availability of nucleotide base.^31,32^ However, the fact that drug-pretreated tumor cells in the presence of nucleosides, a condition able to restore mTORC1 activity, still enhance CD8+ T cell killing suggests that compromised mTORC1 activity in tumor cells is unlikely a reason for enhanced T cell killing. To dissect whether direct mTORC1 inhibition in tumor cells contributes to enhanced CD8+ T cell killing, we employed shRNA-mediated mTOR knockdown or Rapamycin (an mTOR inhibitor) pretreatment. Despite efficient knockdown of mTOR at the protein level and decent reduction in its activity (Figure S5C), tumor cells did not alter the potency of CD8+ T cells as examined by IFN-γ secretion (Figure S5D). In striking contrast, Rapamycin pretreatment of tumor cells dramatically reduced IFN-γ secretion during co-culture (Figure S5E). There is no significant change in tumor antigen presentation after Rapamycin treatment, since membrane expression of HLA-A, B, C is unaltered (Figure S5F) and expression of the tumor antigen NY-ESO-1 is relatively stable, with only subtle reduction when Rapamycin concentration is high (Figure S5G, 200 nM). This makes sense as mTOR inhibition suppresses protein synthesis. Therefore, it is feasible that direct mTOR inhibition only promotes tumor cell resistance to CD8+ T cell killing because it requires antigen processing and presentation (APP) that necessitate mTORC1-governed protein synthesis. The discrepancy observed between different mTOR inhibition strategies may be attributed to different inhibition efficiency of mTOR activity. Nevertheless, our data argue for a conclusion that MTHFD2i- or LTX-enhanced tumor cell sensitivity to T cell killing is not mediated by nucleotide deficiency-resulted reduction in mTORC1 signaling.

After excluding downregulation of mTORC1 signaling as a mechanism for MTHFD2i- or LTX-induced enhancement of tumor immune response, we focused on glycolysis, which is predominantly used by tumor cells for energy production and is consistently downregulated in drug-treated A375 cells (Figure 2A). To explore whether these drugs result in glycolysis suppression, we first analyzed our RNA-seq data concentrating on genes involved in glycolysis. MTHFD2i or LTX treatment significantly downregulated multiple key glycolytic genes, including *HK1, PGAM1*, and *LDHA* (Figure 3A). In agreement, downregulated expression of an important glycolytic enzymes, *LDHA* can be reproduced by polymerase chain reaction (PCR)-based assay in both A375 and HCT 116 cells (Figures 3B and S6A), and this transcriptional suppression is mirrored at the protein level, as shown by decreased abundance of key glycolytic enzymes HK2, PKM2, and LDHA (Figures 3C and S6B). Notably, such downregulation in expression can be translated into functionality, as extracellular lactate production and intracellular ATP levels are dramatically reduced after drug treatment, whereas intracellular glucose levels reversely accumulate (Figures 3D and 3E). These metabolic changes are similarly observed in HCT 116 cells (Figures S6C and S6D). Moreover, genetic suppression of MTHFD2 phenocopied the pharmacological effects, as A375 cells with MTHFD2 knockdown also exhibited reduced lactate production and decreased ATP levels (Figure S6E). Altogether, glucose accumulation coupled with lactate and ATP reduction clearly indicates impaired glycolytic flux, demonstrating that these drugs systematically suppress glycolysis and disrupt cellular energy homeostasis.

**Figure 3.**
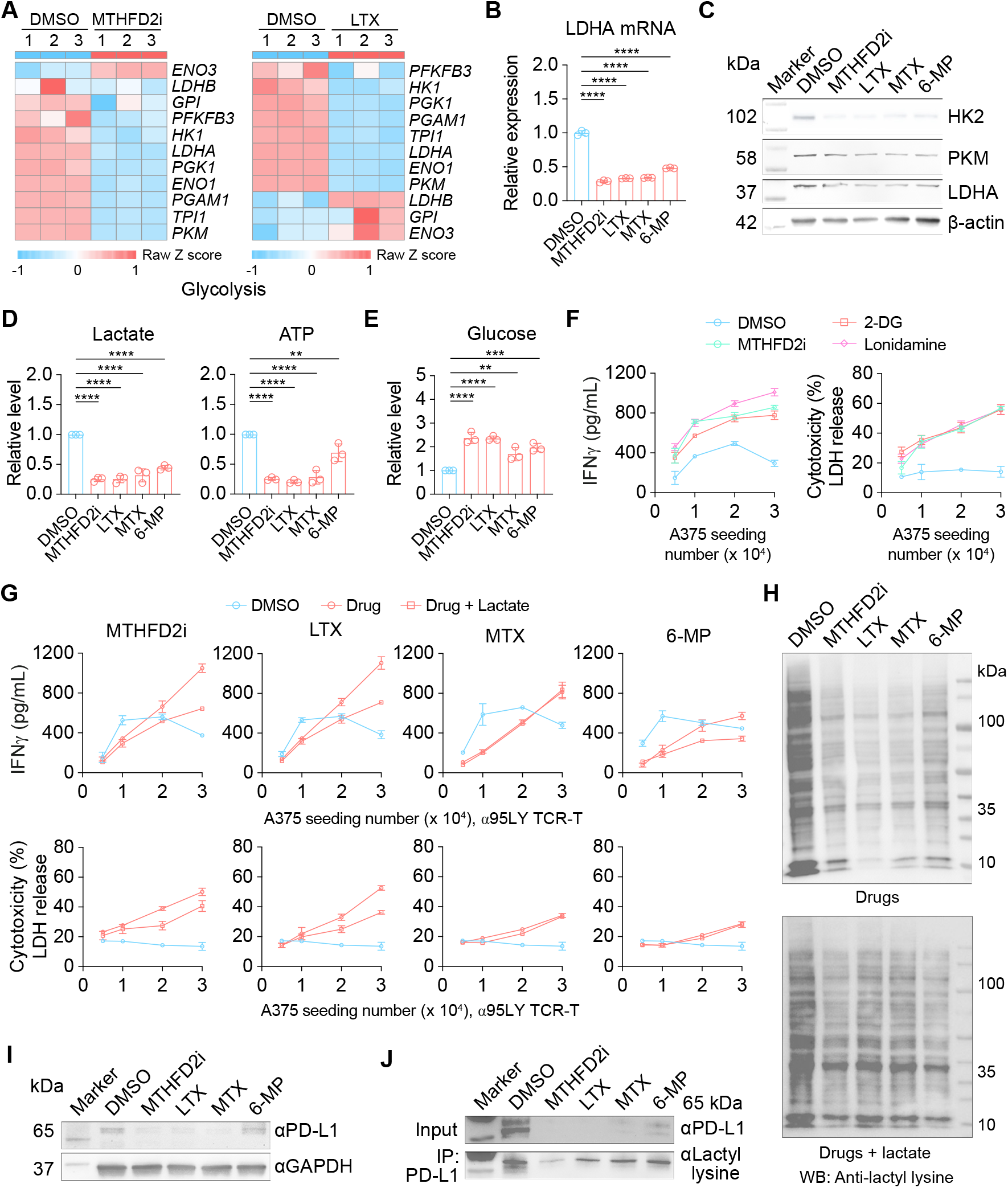
Inhibition of folate-dependent purine synthesis suppresses glycolysis and enhances CD8+ T cell-mediated antitumor immunity. (A) Heatmap showing RNA-seq analysis of glycolysis-related gene expression in A375 cells treated with MTHFD2i (5 µM) or LTX (2 µM) for 24 h. (B) qPCR analysis of LDHA mRNA expression in A375 cells treated with the indicated inhibitors for 24 h. (C) Western blot analysis showing protein levels of HK2, PKM2, and LDHA in A375 cells after treatment with DMSO, MTHFD2i (5 µM), LTX (2 µM), MTX (2 µM), or 6-MP (100 µM) for 24 h. (D, E) Quantification of lactate, ATP, and glucose levels after treatment with DMSO, MTHFD2i (5 µM), LTX (2 µM), MTX (2 µM), or 6-MP (100 µM) for 24 h. (F) IFN-γ secretion in co-culture supernatants. A375 cells seeded at indicated densities were treated with DMSO, MTHFD2i (5 µM), 2-DG (50 µM), or Lonidamine (50 µM) for 24 h, then co-cultured with CD8+ 1G4 α95LY TCR-T cells. (G) IFN-γ secretion (upper panels) and tumor LDH release (lower panels) in co-culture assays. A375 cells seeded at indicated densities were treated with DMSO, MTHFD2i (5 µM), LTX (2 µM), MTX (2 µM), or 6-MP (100 µM) for 24 h, followed by co-culture with CD8+ 1G4 α95LY TCR-T cells in the presence or absence of sodium L-lactate (10 mM). (H) Western blot analysis of global protein lactylation. A375 cells were treated with DMSO, MTHFD2i (5 µM), LTX (2 µM), MTX (2 µM), or 6-MP (100 µM) for 24 h in the absence (Drug) or presence (Drug + lactate) of sodium L-lactate (10 mM) and detected for global protein lactylation. (I) Western blot analysis showing PD-L1 protein levels in A375 cells after treatment with DMSO, MTHFD2i (5 µM), LTX (2 µM), MTX (2 µM), or 6-MP (100 µM) for 24 h. (J) Immunoprecipitation of PD-L1 followed by immunoblotting of PD-L1 lactylation in A375 cells treated with each inhibitor. Data are representative of 3 independent experiments and are presented as mean ± SD. **: p < 0.01; ***: p < 0.001; ****: p < 0.0001.

### Glycolysis-associated lactate reduction mediates the immune response of drug-treated tumor cells

To further explore the link between tumor cell glycolytic status and CD8+ T cell-mediated immune response, we pretreated A375 cells separately with two inhibitors, 2-DG and Lonidamine, both of which target Hexokinase (HK1/2), the enzyme catalyzing the very first step of glycolysis by phosphorylating glucose to produce glucose-6-phosphate. Like MTHFD2i, 2-DG and Lonidamine pretreatment significantly enhanced IFN-γ secretion by CD8+ T cells (Figure 3F). Similar trends were observed in HCT 116 co-culture assays (Figure S6F). These data support the notion that glycolysis inhibition is probably critical in mediating the effect of MTHFD2i or LTX in the context of tumor cell response to T cells.

Lactate is a key metabolite generated by tumor glycolysis. Its accumulation in TME suppresses the function of effector T cells, NK cells, and dendritic cells yet promotes the proliferation and activation of immunosuppressive regulatory T cells (Tregs) and inflammatory (M2) macrophages.^36-38^ Therefore, lactate level is negatively associated with anti-tumor immune response. As is seen reduced when nucleotide synthesis is blocked (Figure 3D), lactate might be a downstream target whose decrease in production could benefit CD8+ T cell killing. To test if this is the case, we added lactate when the nucleotide synthesis drugs were used for tumor cell pretreatment. Indeed, exogenous lactate supplementation attenuated drug-induced enhancement of IFN-γ secretion and tumor cell lysis in our co-culture system (Figures 3G and S6G), highlighting the importance of reduced tumor lactate production in promoting CD8+ T cell responses upon the inhibition of folate-dependent purine synthesis. Of note, MTX-treated tumor cells are insensitive to lactate supplementation, which might be caused by too strong inhibition of tumor cell growth from MTX. Indeed, after drug treatment, lactate supplementation rescued the growth of A375 cells slightly in the case of MTHFD2i or LTX but not MTX (Figure S6H), suggesting persistent inhibition by MTX.

Apart from contributing to an acidic TME, lactate has recently been identified as a donor for lysine residue lactylation, a novel form of post-translational modification occurring on a wide range of proteins, including non-histone proteins.^39,40^ Lactylation of PD-L1, a critical immune checkpoint protein on tumor cells that negatively regulates TCR function, has been shown to contribute to the maintenance of its stability and thereby the enhancement of its immunosuppressive function.^41^ This prompted us to look at protein lactylation after drug treatment. In line with decreased lactate levels, global protein lactylation was reduced when cells were treated with these inhibitors, and lactate supplementation partially restored this modification (Figures 3H and S6I). More specifically, drug treatment not only reduced the overall PD-L1 protein level but also its lactylation (Figures 3I and 3J), suggesting that alterations in lactate metabolism directly impact tumor-immune interactions by regulating PD-L1 lactylation and stability. In summary, our study demonstrates that tumor nucleotide synthesis inhibition decreases PD-L1 stability through attenuation of its lactylation, and meanwhile lactate reduction could potentially alleviate low pH-induced metabolic suppression of CD8+ T cells and ultimately restore tumor sensitivity to immune surveillance.

### Folate-dependent purine synthesis inhibition led to block of ribose salvage

Although we have provided evidence suggesting that impaired glycolysis underlies the enhancement of MTHFD2i- and LTX-induced tumor cell sensitivity to CD8+ T cell killing, the metabolic insight that links impaired folate-dependent purine synthesis and suppressed glycolysis is still lacking. To address this, we performed untargeted metabolome analysis through liquid chromatography with mass spectrometry (LC-MS). Given that MTHFD2i and LTX consistently showed superior sensitizing effects than the other two drugs across our assays, we primarily focused on them in the following studies. Metabolomic samples were prepared after 24-hour treatment of A375 cells with DMSO, MTHFD2i or LTX supplement with nucleosides (cytidine, guanosine, adenosine, uridine, and thymidine), which could theoretically restore the nucleotide pool. Under such condition, any persistent metabolic defects might be directly linked to the enhancement of tumor immune sensitivity (Figure 2C). Our data revealed that MTHFD2i or LTX caused dramatic changes in the abundance of metabolites that are enriched in metabolic pathways, including nucleotide metabolism, purine metabolism, and pyrimidine metabolism (Figure 4A, PathView, KO01232, KO00230, and KO00240), which agrees with our conclusion so far that these drugs predominantly disrupt nucleotide synthesis. Unexpectedly, changes in amino sugar and nucleotide sugar metabolism came to the top. Of note, nucleotide sugar is associated with the hexosamine pathway, which is primarily responsible for protein glycosylation, a post-translational modification with crucial biological functions.^42^ In support, we observed reduction of uridine-derived metabolites, including uridine 5’-triphosphate (UTP) and UDP-N-acetylglucosamine (UDP-GlcNAc) that are metabolites in the hexosamine pathway (Figures 4B and 4C). Additionally, fructose 6-phosphate, a metabolite in the upper hexosamine pathway, is also shown dramatically reduced (Figure 4D). These data point to the view that the hexosamine pathway is significantly compromised, which agrees with previous reports.^43^

**Figure 4.**
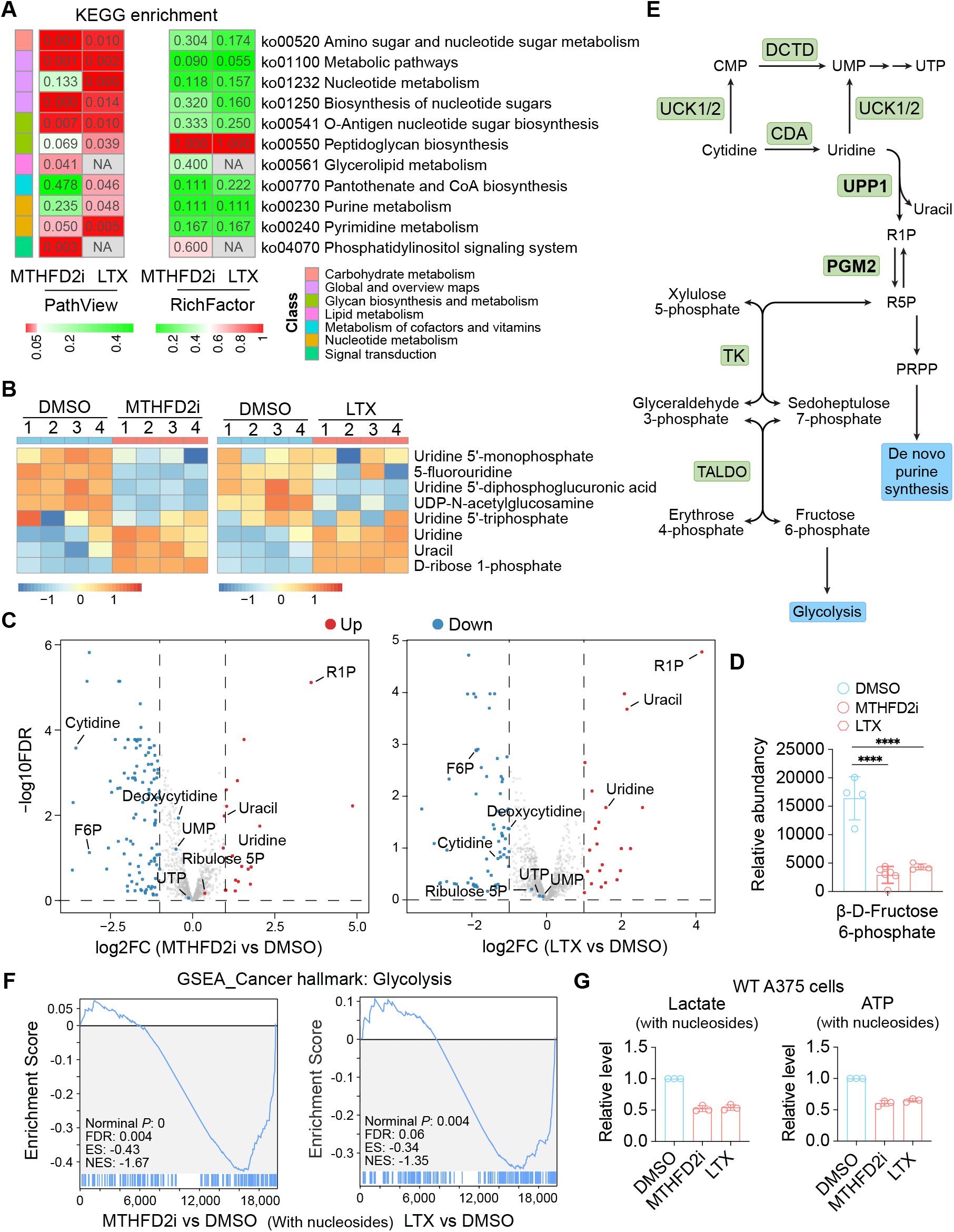
Folate-dependent purine synthesis inhibition alters uridine metabolism and disrupts glycolysis. (A) KEGG pathway enrichment heatmap of differential metabolites in A375 cells treated with DMSO, MTHFD2i (5 μM) or LTX (2 μM) for 24 h with nucleoside supplementation. Nucleosides were added in a final concentration of 30 μM for cytidine, guanosine, adenosine, and uridine each, and 10 μM for thymidine. Adjusted P values (FDR, left) for pathway enrichment and RichFactor (right) are shown. (B) Heatmap of the metabolomic data showing changes in uridine-related metabolites. (C) Volcano plots of the metabolomic data. Metabolites with more than 2-fold change are highlighted and representative metabolites are annotated. (D) Relative abundance of β-D-Fructose-6-phosphate from metabolomic data. ****: p < 0.0001. (E) Schematic diagram illustrating the metabolic relationship of uridine metabolism and glycolysis, highlighting the role of UPP1 and PGM2 in linking these metabolic pathways. (F) GSEA plots showing downregulation of glycolysis-related gene sets in A375 cells treated with MTHFD2i (5 µM) or LTX (2 µM) for 24 h versus DMSO. (G) Quantification of lactate and ATP levels in A375 cells treated with DMSO, MTHFD2i (5 µM), or LTX (2 µM) for 24 h in the presence of nucleosides (30 μM for cytidine, guanosine, adenosine, and uridine each, and 10 μM for thymidine).

Since the hexosamine pathway is heavily reliant on uridine metabolism, its downregulation might be an indication of abnormalities in uridine metabolism. Indeed, we observed an elevation of uridine phosphorylytic cleavage products uracil and D-ribose 1-phosphate (R1P) (Figures 4B and 4C). These data provide clue of the occurrence of disruption in uridine metabolism, which favors uridine phosphorylytic cleavage at the sacrifice of nucleotide or nucleotide sugar production. However, UMP and UTP levels are only mildly reduced, though pathway analysis, together with the observation of abnormal uridine accumulation, suggests uridine-derived nucleotide synthesis might be blocked. Strikingly, we found that cytidine level is drastically lowered in drug-treated groups (Figure 4C), indicating it might be used as an alternative branch for UMP synthesis (Figure 4E).

When nutrient is limited, uridine serves as an essential source of energy material and provides ribose to fuel glycolysis for energy production, a pathway termed ribose salvage and adapted by a wide range of tumor cell lines under energy stress.^13,14^ Downstream, R1P obtained from uridine is converted into Ribose 5-phosphate (R5P) before the latter is used for the production of two glycolytic intermediates glyceraldehyde 3-phosphate and fructose 6-phosphate through non-oxPPP (Figure 4E).^11^ For unknown reason, R1P is prevented from entering the ribose salvage pathway and therefore is excessively accumulated (Figures 4B and 4C). Consequently, glycolysis is under pressure even in the presence of nucleosides, as revealed by reduced fructose 6-phosphate (Figure 4D), downregulated glycolysis from GSEA analysis (Figure 4F), and decreased cellular lactate and ATP levels (Figure 4G). Taken together, although nucleoside supplementation can rescue DNA synthesis and mTOR activity, MTHFD2i or LTX applies persistent stress on nutrient consumption pathways involving nucleotide sugar metabolism, which appears to be correlated with block of ribose salvage.

### UPP1 interferes with drug-induced suppression of glycolysis and enhancement of tumor immune sensitivity

Our metabolome analysis directed us to look at key enzymes that catalyze the conversion of uridine into uracil and R1P, and the conversion of R1P into R5P that can be utilized for glycolysis through ribose salvage (Figure 4E). The accumulation of R1P suggests the activity of these enzymes might be influenced by drugs, and therefore it is intriguing to test whether manipulating the expression of these enzymes can recapitulate drug effects. Uridine phosphorylase 1 (UPP1) is responsible for catalyzing the reversible phosphorylation of uridine, cleaving uridine into uracil and R1P.^44^ UPP1 is found essential to tumor cells through providing ribose for glycolysis during energy stress.^13,14,45,46^ Evidenced by the accumulation of uracil and R1P in MTHFD2i- or LTX-treated A375 cells supplemented with nucleosides (Figures 4B and 4C), we anticipated an active participation of UPP1 in such scenario. However, UPP1 transcripts in A375 cells were consistently reduced after MTHFD2i or LTX treatment (Figure 5A), indicating folate-dependent purine synthesis inhibition suppresses glycolysis partially through downregulating UPP1 expression. This notion is confirmed when we measure glycolytic gene expression and perform functional assays in UPP1-overexpressing A375 cells treated with MTHFD2i or LTX. UPP1 overexpression brought about strong recovery of HK2 and LDHA mRNA levels (Figure 5B) as well as their protein levels (Figure 5C). As a result, lactate reduction was also mostly restored (Figure 5D). Functionally, UPP1 overexpression not only reduced tumor cell susceptibility, but also compromised MTHFD2i- or LTX-induced enhancement of tumor cell sensitivity to CD8+ T cell killing, as reflected by decreased IFN-γ secretion and reduced LDH release in the co-culture assays (Figure 5E). These data demonstrate that with respect to glycolysis, UPP1 plays an opposing role against MTHFD2i and LTX.

**Figure 5.**
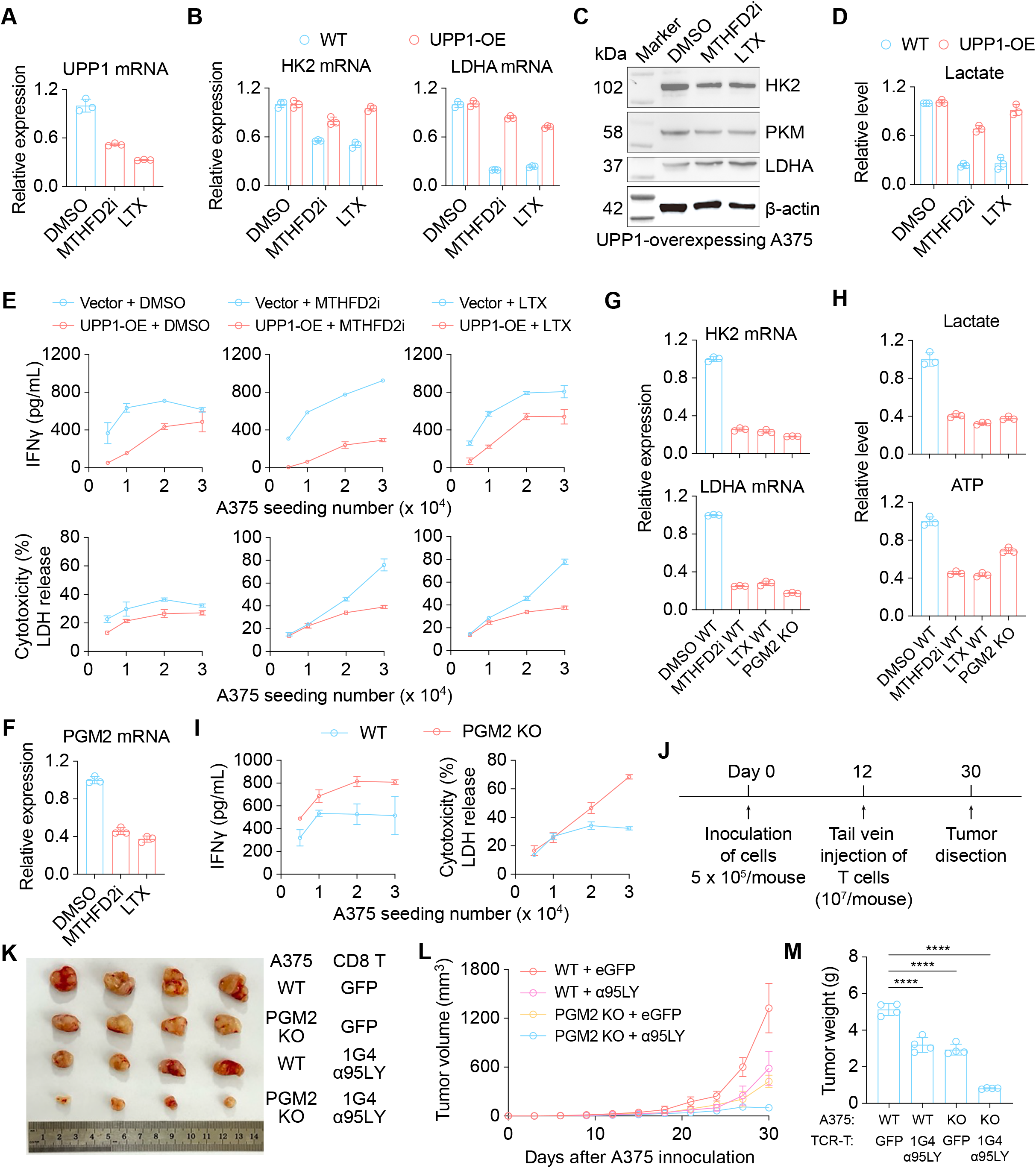
UPP1 and PGM2 interferes with glycolysis and enhancement of tumor immune response. (A) qPCR analysis of UPP1 mRNA levels in A375 cells treated with DMSO, MTHFD2i (5 µM), or LTX (2 µM) for 24 h. (B) qPCR analysis of HK2 and LDHA mRNA levels in WT and UPP1-overexpressing (UPP1-OE) A375 cells treated with DMSO, MTHFD2i (5 µM), or LTX (2 µM) for 24 h. (C) Western blot analysis of HK2, PKM2, and LDHA protein levels in A375 cells treated with DMSO, MTHFD2i (5 µM), or LTX (2 µM) for 24 h. (D) Lactate quantification in WT and UPP1-OE A375 cells treated with DMSO, MTHFD2i (5 µM), or LTX (2 µM) for 24 h. (E) IFN-γ secretion and tumor LDH release in co-culture assays. WT and UPP1-OE A375 cells seeded at indicated densities were treated with DMSO, MTHFD2i (5 µM), or LTX (2 µM) for 24 h and then co-cultured with CD8+ 1G4 α95LY TCR-T cells. (F) qPCR analysis of PGM2 mRNA levels in A375 cells treated with DMSO, MTHFD2i (5 µM), or LTX (2 µM) for 24 h. (G) qPCR analysis of HK2 and LDHA mRNA levels in WT A375 cells treated with DMSO, MTHFD2i (5 µM), or LTX (2 µM) for 24 h, as well as those in PGM2 knockout (PGM2 KO) A375 cells. (H) Relative quantification of lactate production and ATP levels in WT A375 cells treated with DMSO, MTHFD2i (5 µM), or LTX (2 µM) for 24 h, and in PGM2-KO A375 cells. (I) IFN-γ secretion and tumor LDH release in co-culture assays. WT and PGM2 KO A375 cells seeded at indicated densities were co-cultured with CD8+ 1G4 α95LY TCR-T cells for 24 h. (J) Schematics of in vivo tumor killing assay. (K) Representative images of excised tumors from the indicated groups at the endpoint. (L) Tumor growth curves of mice bearing WT or PGM2 KO A375 tumors after adoptive transfer of control eGFP- or 1G4 α95LY-transduced CD8+ T cells. (M) Tumor weights of the indicated groups at the endpoint. Each dot represents one mouse. Statistics were calculated with t-test. ****: *P* < 0.0001.

### Loss of PGM2 suppresses glycolysis and sensitizes melanoma cells to CD8+ T cell killing

The observed accumulation R1P can hardly be explained by the downregulation of UPP1, which would otherwise lead to reduced R1P. We thus hypothesized that the activity of phosphoglucomutase 2 (PGM2), the enzyme critical for the conversion of R1P into R5P to fuel glycolysis, is severely compromised. Indeed, PGM2 expression is also dramatically reduced after drug treatment (Figure 5F). To examine its involvement in glycolysis, we performed CRISPR/Cas9-mediated knockout of PGM2 in A375 cells. As a result, deletion of PGM2 reduced the expression HK2 and LDHA (Figure 5G), as well as intracellular lactate and ATP levels comparable to drug effects (Figure 5H), consistent with PGM2 serving as a functional bridge linking uridine-derived ribose to glycolysis. Importantly, PGM2 deletion in tumor cells promoted T cell effector responses, as evidenced by increased IFN-γ secretion and enhanced tumor cell lysis (Figure 5I). Moreover, in NOG (*NOD*.*Cg-Prkdc*^*scid*^*Il2rg*^*tm1Sug*^*/JicCrl*) immune-deficient mouse xenograft model (Figure 5J), tumor growth from PGM2-deficient A375 cells is dramatically inhibited in the presence of tail-vein injected 1G4 α95LY TCR-T cells (Figures 5K-5M). Taken together, the observation of uracil and R1P accumulation might be a sign of functional impairment of PGM2, which blocks the entry of R1P into glycolysis via non-oxPPP and damages glycolysis. Loss of PGM2 significantly improves TCR-T cell-mediated tumor immune attack in immune-deficient mouse model.

### Drug impairs uridine metabolism and enhances tumor immunity through downregulating UTP-sensitive genes

In addition to the accumulation of uracil and R1P, MTHFD2i or LTX treatment also led to decrease in uridine derivatives (Figure 4B), indicating that these drugs strongly impair uridine metabolism such that supplemented uridine is neither utilized as an alternative nutrient nor efficiently devoted to the generation of nucleotide sugars for protein modification or materials for RNA synthesis. This led us to ask whether such defect would influence transcription and gene expression that could potentially sensitize tumor cells for T cell immunity. Therefore, we performed RNA-seq of MTHFD2i- or LTX-treated A375 cells supplemented with nucleosides. Along with the recovery of DNA synthesis and mTOR activity, the expression of drug-induced differentially expressed genes mostly returned to normal when nucleosides were supplied (Figure 6A). However, there were a few genes showing resilience in restoring expression, which are a group of heat shock proteins functionally involved in protein folding (Figures 6A and 6B). These findings can be reproduced by PCR quantification (Figure 6C). Since uridine metabolism is still defective in drug-treated cells with nucleoside supplementation, showing shortage in uridine derivatives including UTP, we argued whether the transcripts of *HSPA1A* and *HSPA1B* are sensitive to uridine metabolism. Notably, after analyzing the 3’ untranslated regions (UTRs) of the human transcriptome, we found significant enrichment of nucleotide U in the 3’ UTR of both transcripts, constituting more than 40% (Figure 6D). Strikingly, the enrichment of nucleotide U in the 3’ UTRs of *HSPA1A* and *HSPA1B* ranks the top of all 3’ UTRs in transcripts, which might account for their sensitivity to drug-induced impairment in uridine metabolism, with reduced uridine derivatives compromising the mRNA levels of these genes. Finally, such downregulation could be translated into functionality as pharmacologically inhibiting tumor HSP70 increased their susceptibility to CD8+ T cell killing (Figure 6E). Altogether, MTHFD2i or LTX enhances tumor immune sensitivity via disrupting uridine metabolism, causing UTP shortage, and reducing the transcript of genes with a protective role against T cells.

**Figure 6.**
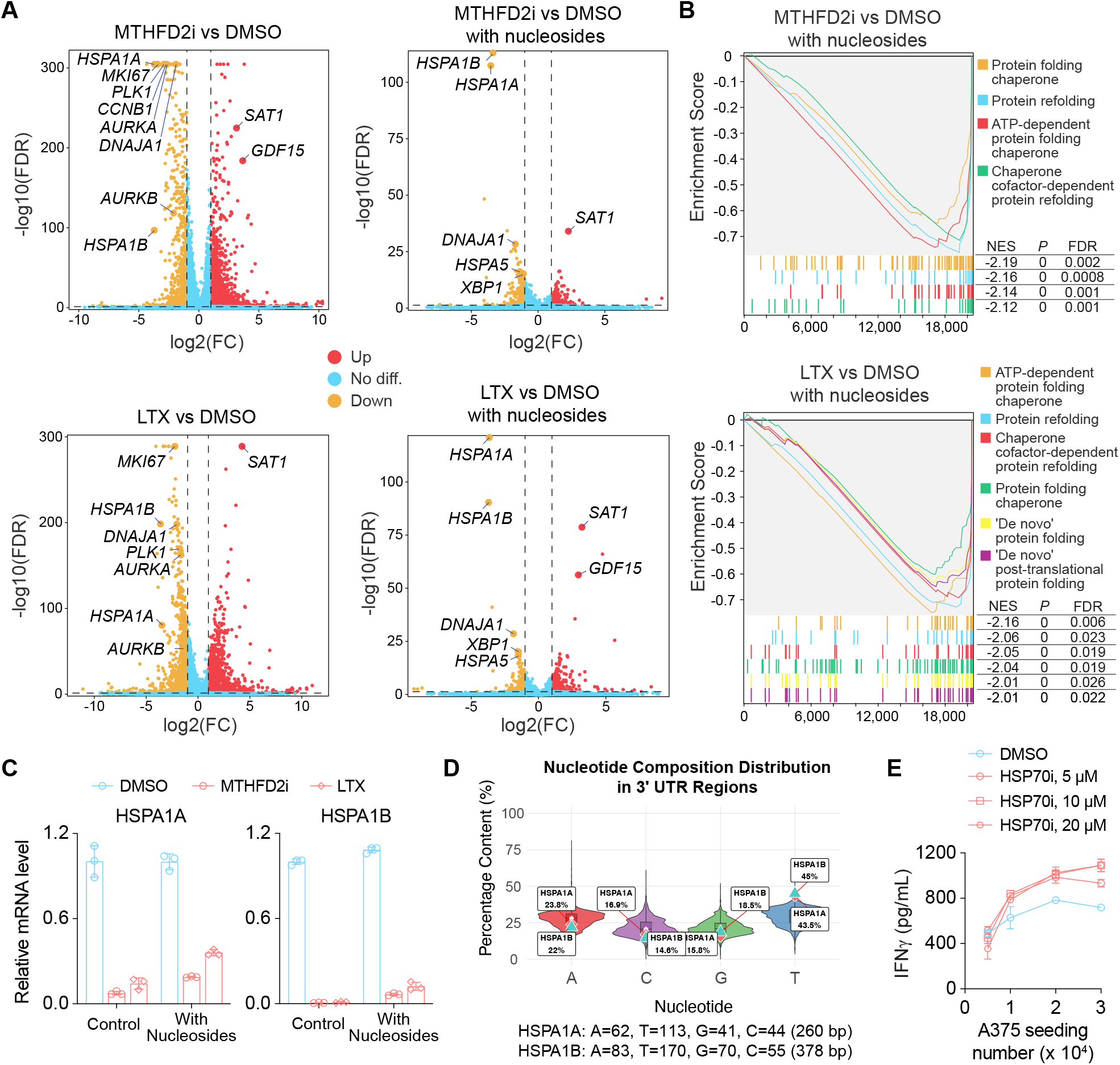
Folate-dependent purine synthesis inhibition downregulates the transcripts of protein folding chaperones. (A) Volcano plot showing differentially expressed genes after treatment with MTHFD2i (5 µM) or LTX (2 µM) versus DMSO with or without nucleoside supplementation. (B) GSEA analysis showing downregulated pathways after nucleoside rescue in drug-treated cells. (C) qPCR analysis of HSPA1A and HSPA1B mRNA levels in A375 cells treated with DMSO, MTHFD2i (5 µM), or LTX (2 µM) with or without nucleoside supplementation for 24 h. (D) Violin plots showing the distribution of A, T, G, C nucleotide frequencies within the 3’ UTRs of HSPA1A and HSPA1B compared to all other genes in the transcriptome. (E) IFN-γ secretion after co-culture. A375 cells seeded at indicated densities were treated with different concentrations of HSP70 inhibitor for 24 h and then co-cultured with CD8+ 1G4 α95LY TCR-T cells.

## Discussion

One-carbon (1C) metabolism cooperates with de novo purine synthesis to support elevated nucleotide demand from proliferating tumor cells.^4^ Drugs targeting these pathways have long been used in the treatment of cancer. Although their role in cell cycle inhibition is well known, how these drugs modulate tumor-immune interaction is relatively unclear. Since cellular metabolism is extremely complex with many pathways interconnected, disturbance on a single molecule would induce widespread metabolic changes. In the present study, we propose that inhibition of MTHFD2 by MTHFD2i or GART by LTX induces suppression of glycolysis, which is a major driver for enhanced tumor cell sensitivity to immune attack. Our findings provide a rationale for leveraging folate-dependent purine synthesis pathway inhibition to enhance tumor sensitivity to T cell-mediated killing.

Glucose is converted to glucose-6-phosphate, which can enter oxPPP to generate R5P for nucleotide synthesis.^11,47^ Conversely, nucleosides/nucleotides undergoes cleavage to produce bases and R1P, which can be converted into R5P to generate glycolytic intermediates (such as glyceraldehyde 3-phosphate and fructose 6-phosphate, Figure 4E) through non-oxPPP.^14,48^ Thus, folate-dependent purine synthesis and glycolysis are metabolically connected through the R1P/R5P node. Our study provides evidence that MTHFD2i or LTX perturbs the coupling between glycolysis and nucleotide metabolism, with uridine metabolism persistently compromised even nucleotide pool is refilled with nucleoside supplementation. The metabolic mechanisms that transduce drug inhibition of purine synthesis to compromised uridine metabolism is still unknown. We speculate that the activity of critical enzymes within this pathway might be altered by drug-induced change in the concentration of key metabolic intermediates, just like the case with transketolase (TK) in non-oxPPP. Specifically, PRPP and adenosine diphosphate (ADP) has been shown to inhibit the activity of TK,^49^ therefore accumulation of PRPP sends negative feedback towards non-oxPPP and restricts ribose to be consumed as fuel. In our case, compromised uridine metabolism is manifested by the accumulation of uridine, uracil, and R1P. To interpret such results, drug-induced alterations in salvage or transport efficiencies of uridine could be responsible. However, our findings that both UPP1 (governing uridine phosphorylytic cleavage) and PGM2 (governing ribose conversion from R1P to R5P) can interfere with glycolysis and tumor immune response suggest alterations in the activities of enzymes catalyzing uridine metabolism should be accountable. Limitations exist that only functional tests were performed with these enzymes without quantification of changes in metabolites. This can be tested in future work using stable-isotope tracing to quantify carbon flux between metabolic pathways.^50,51^ Nevertheless, we offer potential new targets, such as PGM2, for cancer treatment.

Additionally, folate-dependent purine synthesis inhibition by MTHFD2i or LTX is associated with reduced expression of key glycolytic regulators, including LDHA, UPP1, and PGM2, which may further contribute to diminished glycolytic activity. These are alternative mechanisms that contribute to reduced glycolytic activity. Glycolysis serves as the predominant energy supply pathway for tumor cells and plays a central role in sustaining an immunosuppressive tumor microenvironment. Mounting evidence indicates that the accumulation of lactate from glycolysis impairs effector T cell function and promotes tumor cell immune evasion. Our work revealed that folate-dependent purine synthesis inhibition suppresses glycolysis and thereby reduces lactate production and PD-L1 lactylation, offering a mechanism on how these inhibitors increase tumor cell susceptibility to CD8+ T cell killing and providing a solution of targeting glycolysis for cancer treatment.

Our experimental design adopted a drug-pretreatment strategy, followed by tumor antigen-specific TCR-T cell killing. This aligns with the idea of combining chemotherapy with adoptive T cell transfer therapy for cancer treatment. Although T cells are avoided to be affected by drugs, the implication of our findings would be wider if drugs do not inhibit the function of T cells. In fact, it has been reported that MTHFD2i impairs CD4+ T cell proliferation and function.^52^ Much like what we have found in tumor cells, it dampens mTORC1 activity and shifts the metabolic program from glycolysis toward oxidative phosphorylation in both Th17 and Treg cells. Phenotypically, MTHFD2i promotes FoxP3 and Treg cell-like phenotypes in Th17 cells, suggesting MTHFD2 may function as a metabolic checkpoint in the Th17-Treg cell axis, with MTHFD2i skewing the balance from a pathogenic to a more anti-inflammatory phenotype. It is interesting that a metabolic intermediate in the de novo purine synthesis pathway, 5-Aminoimidazole-4-carboxamide ribonucleotide (AICAR), is accumulated in MTHFD2i-treated CD4 T cells. AICAR acts as an indirect agonist for AMPK, which suppresses mTORC1 activity. This might explain why MTHFD2i inhibits mTORC1 activity in CD4+ T cells. However, our data revealed downregulation of AICAR in MTHFD2i-treated A375 cells with nucleoside supplementation (Figure S7), which helps to interpret the data showing restored mTORC1 activity after nucleoside supplementation, since downregulation of AICAR lifts AMPK suppression of mTORC1. These contrasting results are probably caused by different experimental settings. Nevertheless, it appears that drugs have significant impact on CD4+ T cells. It is unknown how drugs influences CD8+ T cells, especially their effector function. Immune-deficient mouse models involving human-derived tumor cell lines, or immune-competent mouse models involving syngeneic mouse tumor cell lines should help to address this question. Future studies should define the generalizability of this metabolic-immune coupling across tumor contexts with distinct reliance on de novo synthesis versus salvage pathways and delineate direct effects of these interventions on T cell fitness and effector function to guide rational combinations with adoptive cell therapies such as TCR-T or CAR-T therapy.

## Conflict of interest

The authors declare no conflict of interest.

## Data availability

RNA-seq datasets generated and analyzed during the current study are available in the Gene Expression Omnibus (GEO) repository with the identification number GSE312437. Currently reviewers can access the data with the secure token utehqcqmtjshxmn.

The untargeted metabolomic dataset is deposited at MetaboLights and can be accessed through https://www.ebi.ac.uk/metabolights/MTBLS13434. Currently it is available to reviewers through the read-only private link https://www.ebi.ac.uk/metabolights/reviewercbee5d56-2eec-4435-9665-15e63e58d700.

## Funding

This work is supported by grants from the National Natural Science Foundation of China (82273498), and the Non-profit Central Research Institute Fund of Chinese Academy of Medical Sciences (2021-RC320-002).

## Methods

### Animals and ethical statement

The NOG (NOD.Cg-PrkdcscidIl2rgtm1Sug/JicCrl) mice used in this study were obtained from Charles River. Mouse work was approved by the Institutional Animal Care and Use Committee of Institute of Dermatology, Hospital for Skin Diseases, Chinese Academy of Medical Sciences & Peking Union Medical College (No. 2023-DW-021). Related experiments were performed in accordance with the ARRIVE guidelines for animal research (https://arriveguidelines.org), ensuring transparency and reproducibility of using animal-derived materials. Six-to eight-week-old mice were used for tumor inoculation. On day 0, A375 cells (5 × 10^5^ cells per mouse) were subcutaneously inoculated into the lateral abdomen. Tumor growth was monitored every 3 days throughout the experiment. On day 12, 1G4 α95LY TCR-T cells were resuspended in RPMI-1640 medium and transferred into anaesthetized mice through tail vein injection, with 1 × 10^7^ cells per mouse in a volume of 120 μL. On day 30, mice were euthanized and tumor were dissected.

### Antibodies

Antibodies against mTOR (7C10, no. 2983S, 1:1,000), phospho-p70 S6 kinase (Thr389) (108D2, no. 9234S, 1:1,000), p70 S6 kinase (49D7, no. 2708S, 1:1,000), HK2 (C64G5, no. 2867S, 1:1,000), PKM2 (D78A4, no. 4053S, 1:1,000), β-actin (13E5, no. 4970S, 1:1,000), and LDHA (C4B5, no. 3582S, 1:1,000) were purchased from Cell Signaling Technology. Anti-HLA-A, B, C (311418, 1:100) was purchased from BioLegend. PD-L1 antibody (ab121009, 1:1,000) was obtained from Aladdin. L-lactyl lysine rabbit polyclonal antibody (S0B0719, 1:1,000) was purchased from Starter Bio. MTHFD2 antibody (ab151447, 1:1,000) was purchased from Abcam.

### Reagents

DS18561882 (HY-130251), lometrexol (HY-14521), methotrexate (HY-14519), 6-mercaptopurine (HY-13677), 2-deoxy-D-glucose (HY-13966), lonidamine (HY-B0486), adenine (HY-B0152), L-Lactic acid sodium (HY-W040233) and the targeted diversity library (HY-L099) were obtained from MedChemExpress (MCE). Rapamycin (V900930-1MG) and EmbryoMax® nucleosides (ES-008-D) were purchased from Sigma-Aldrich.

### Cell lines and culture conditions

A375, HCT 116, A498, and 293FT cells were obtained from the National Collection of Authenticated Cell Cultures at the Institute of Biochemistry and Cell Biology, Chinese Academy of Sciences (Shanghai, China). A375, 293FT cells were cultured in DMEM medium (Gibco, C11995500BT) supplemented with 10% fetal bovine serum (FBS), 1× penicillin/streptomycin (Gibco, 15140122), and 1× GlutaMAX™ (Gibco, 35050061). A498 cells were cultured in α-MEM (Gibco, 12571063) supplemented with 10% FBS, 1× penicillin/streptomycin, and 1× GlutaMAX™. HCT 116 cells were maintained in McCoy’s 5A medium (Gibco, 16600082) supplemented with 10% FBS, 1× penicillin/streptomycin, and 1× GlutaMAX™. Human primary T cells were isolated from peripheral blood mononuclear cells (PBMCs) obtained from a local blood center and cultured in AIM-V medium (Gibco, 12055083) supplemented with 10% human AB serum (GemCell, 100-512), 1× penicillin/streptomycin, 1× GlutaMAX™, and 300 U/mL recombinant human IL-2 (Peprotech, 200-02-100). All cells were maintained at 37 °C in a humidified atmosphere with 5% CO_2_.

### Plasmids, lentiviral packaging, and viral transduction

Plasmids encoding 1G4 α95LY TCR and MAGE-A3 94a-14 TCR were described previously.^18^ UPP1 coding sequence was obtained from Miaoling Bio. (P74511) and subcloned into a lab-customed lentiviral vector pLV3-EF1α-2A-puro. Lentiviral particles were produced in 293FT cells by co-transfection with DR8.91 and VSVG plasmids. Retroviral particles were generated in 293FT cells by co-transfection with pGP (Takara) and RD114 plasmids. Lentiviral transduction was performed in the presence of 5 μg/mL polybrene (Millipore, TR-1003-G). For retroviral transduction of human primary CD8+ T cells, PBMCs were isolated using Lymphoprep™ (Stem Cell Technologies, 07851), and CD8+ T cells were purified using the EasySep™ Human CD8+ T Cell Isolation Kit (Stem Cell Technologies, 17953). Purified CD8+ T cells were activated by Dynabeads™ Human T⍰Activator CD3/CD28 (Gibco, 11161D) at a 1:1 bead⍰to⍰cell ratio in round⍰bottom 96⍰well plates for 36 – 48 h. Retroviral transduction was carried out on RetroNectin (Takara, TKR⍰T100A)⍰coated plates following the manufacturer’s instructions. Briefly, retroviral supernatant was centrifuged onto the RetroNectin⍰coated plates. Activated T cells were then detached from the Dynabeads, transferred to the virus⍰coated plates, and centrifuged at 2,000 rpm for 2 h at 32°C before being incubated overnight.

### Co-culture of T cell killing

For co-culture with single density of tumor cells, 1 × 10^4^ cells were seeded in one well of 96-well plates with 3 replicates for each condition. For co-culture with multiple densities of tumor cells, 0.5 ×, 1 ×, 2 ×, and 3 × 10^4^ cells were seeded into one well of 96-well plates, respectively, with 3 replicates for each condition. Cells were allowed to grow for 24 hours before treatments were performed. Drugs with indicated concentrations were used to treat tumor cells for 24 hours before 1 × 10^4^ TCR-T cells with at least 70% positivity were added. In case of no drug treatment, 1 × 10^4^ TCR-T cells were added 24 hours after seeding of tumor cells. Tumor – T cell co-culture was performed for another 24 hours before supernatants were collected for Elisa assay.

### Construction of knockdown cell lines

Doxycycline-inducible mTOR knockdown A375 cell lines were generated using a TetO-based lentiviral shRNA system (pLKO-puro-TetO vector). Stable MTHFD2 knockdown A375 cell lines were generated using a lentiviral shRNA system (pLKO.1-mCherry-Puro vector, Miaoling Bio., P10494). For construction of shRNA plasmids, complementary oligonucleotides were annealed and cloned into corresponding vectors. All constructs were verified by Sanger sequencing. Lentiviral particles were produced and used to transduce A375 cells. 48 h post transduction, cells were selected with puromycin (2 μg/mL) overnight to eliminate non-expressers. For mTOR, knockdown was induced with doxycycline (1 μg/mL). Knockdown efficiency was assessed by immunoblotting. Targeted sequences are listed as follows:

shMTHFD2-1: CGAGAAGTGCTGAAGTCTAAA

shMTHFD2-2: GCAGTTGAAGAAACATACAAT

shmTOR-1: CCGCATTGTCTCTATCAAGTT

shmTOR-2: TCAGCGTCCCTACCTTCTTCT

shTurboGFP: CGTGATCTTCACCGACAAGAT

### Construction of PGM2 knockout A375 single cell clones

PGM2 knockout A375 cells were generated using CRISPR/Cas9 technology. Two sgRNAs targeting PGM2 were separately cloned into a lab-customed CRISPR/Cas9 plasmid pLenti-CRISPR-puro-V3. A375 cells were transfected with targeting plasmids, expanded for 48 hours, and selected with puromycin (2 μg/mL) for 24 hours. Cells were allowed to recover for 3 – 4 days before single cells were seeded by limiting dilution into 96-well plates covered with feeder layer, with a theoretical density of one cell per well. Typically, after 1 – 2 weeks, wells with single colonies were selected and expanded. Successful knockout was verified by genotyping.

sgPGM2-1: GTGAAACGACTAATAGCAGA

sgPGM2-2: GAGCTCATCCATCCAGTGGG

### Enzyme-linked immunosorbent assay (ELISA)

Supernatants were collected from co-cultures of tumor cells and TCR-T cells. ELISA reagents were purchased from eBioscience (88-7316-77). Briefly, ELISA plates were coated with 100 μL of capture antibody in coating buffer and incubated overnight at 4°C. Plates were blocked with 200 μL of ELISA diluent for 1 hour at room temperature before 100 μL of sample or standard solutions were added to the plates and incubated overnight at 4°C. After removal of the samples, 100 μL of TMB substrate was added to develop color for 15 minutes. Color development was stopped by stop solution and absorbance was recorded at 450 nm or 470 nm.

### Lactate dehydrogenase (LDH) release Assay

LDH release was quantified according to the manufacturer’s instructions (Beyotime, C0017). Specifically, cells were seeded in 96-well plates and measurement of LDH activity in the culture supernatants was performed when cells reach confluency of 80 – 90%. For experimental design, four groups of treatments were included as follows: medium-only group as background control, cells untreated as spontaneous release control, cells untreated but subject to lysis before quantification as maximum release control, and cells treated with indicated drugs including vehicle control. One hour before measurement, 10% of the culture volume of LDH release reagent was added to the group of maximum release control. Cells were incubated to the endpoint. Plates were centrifuged at 400 × g for 5 min and 120 μL of the supernatant from each well was transferred to a new 96-well plate for measurement of LDH activity.

### Live cell imaging

A375 cells were seeded into 96-well plates at 1 × 10^4^ cells/well. 24 hours later, cells were refreshed with medium containing DMSO or MTHFD2i (5 µM) and incubated for another 24 hours before human primary T cells expressing GFP and 1G4 α95LY TCRs were added at a 1:1 effector-to-target (E:T) ratio. Images were captured using the Incucyte® Live-Cell Imaging System (Sartorius).

### Cell viability assay (CCK-8)

Cell viability was measured using Cell Counting Kit-8 (CCK-8, Vazyme, A311-01). Cells were seeded in 96-well plates and treated as indicated. CCK-8 reagent (10 μL) was added to each well containing 100 μL medium and incubated for 1 h at 37°C protected from light. Absorbance was read at 450 nm using a microplate reader. Cell viability was normalized to the corresponding control after background subtraction. Each condition was tested in technical replicates, and experiments were repeated three times independently.

### RNA-seq and data analysis

Total RNA was extracted using Trizol reagent (Invitrogen, 15596-026) following the manufacturer’s instructions. RNA quality was assessed by RNase-free agarose gel electrophoresis and an Agilent 2100 Bioanalyzer. mRNA was enriched using Oligo(dT) beads, fragmented, and reverse transcribed into cDNA with the NEBNext Ultra RNA Library Prep Kit for Illumina (New England Biolabs, E7770). The resulting cDNA libraries were sequenced on an Illumina NovaSeq 6000 platform by Gene Denovo Biotechnology Co. (Guangzhou, China). Sequencing reads passing quality control were first aligned with Bowtie2 (version 2.2.8) to remove rRNA-mapped short reads. Clean reads were then mapped to the human reference genome (GRCh38) using HISAT2 (version 2.2.4) with default parameters. Transcript assembly was performed in a reference-guided manner using StringTie (version 1.3.1). Transcript abundance was quantified as FPKM (fragments per kilobase of transcript per million mapped reads) values using RSEM. Differentially expressed genes (DEGs) between groups were identified with DESeq2, applying a false discovery rate (FDR) < 0.05 and an absolute fold change ≥ 2 as the significance threshold. Gene Set Enrichment Analysis (GSEA) was conducted with GSEA software and MSigDB, using the gene expression matrix as input and Signal-to-Noise normalization to rank genes. Enrichment scores and p values were obtained with default settings. Gene expression heatmaps were generated in R.

### Reverse transcription-quantitative polymerase chain reaction (RT-qPCR)

Total RNA from cells was extracted using TRIzol reagent (Invitrogen, 15596026) and reverse transcribed into cDNA with the HiScript III 1st Strand cDNA Synthesis Kit (Vazyme, R312-01). Quantitative real-time PCR was performed using AceQ® qPCR SYBR Green Master Mix (Vazyme, Q511). Relative gene expression levels were calculated using the 2^−ΔΔCt method and normalized to GAPDH. Primer sequences used for qPCR are listed as follows.

UPP1-F: TGATTGCCCCGTCAGACTTTT

UPP1-R: CACCAACGCACCTGATGAAG

HK2-F: GAGCCACCACTCACCCTACT

HK2-R: CCAGGCATTCGGCAATGTG

PKM2-F: ATGTCGAAGCCCCATAGTGAA

PKM2-R: TGGGTGGTGAATCAATGTCCA

LDHA-F: ATGGCAACTCTAAAGGATCAGC

LDHA-R: CCAACCCCAACAACTGTAATCT

HSPA1A-F: GGTGCTGACGAAGATCAAGGAGATC

HSPA1A-R: CTGCCGCTGACAGTCGTTGAAG

HSPA1B-F: GCCTTTCCAAGATTGCTGTT

HSPA1B-R: TCAACATTGCAAACACAGGA

### Immunoprecipitation and Western Blotting

Total protein was extracted from cultured cells using ice-cold Pierce IP lysis buffer (Thermo Scientific, 87787) supplemented with protease and phosphatase inhibitor cocktails. For immunoprecipitation, cell lysates were incubated overnight at 4 °C with PD-L1 antibody, followed by protein pull-down and immunoblotting with the pan-lactylation antibody. For Western blotting, equal amounts of protein were separated by SDS-PAGE and transferred onto PVDF membranes (Bio-Rad). The membranes were then probed with the following primary antibodies: PD-L1 (Aladdin, 1:1000), pan-lactylation (Starter, 1:1000), mTOR (7C10, #2983S, 1:1000), Phospho-p70 S6 Kinase (Thr389) (108D2, #9234S, 1:1000), p70 S6 Kinase (49D7, #2708S, 1:1000), HK2 (C64G5, #2867, 1:1000), PKM2 (D78A4, #4053, 1:1000), β-Actin (13E5, #4970S, 1:1000), and LDHA (C4B5, #3582, 1:1000). Protein bands were visualized using a chemiluminescence imaging system (Bio-Rad).

### Flow cytometry

Cell cycle was analyzed using the FUCCI Amcyan-hGem/mCherry-hCdt1 system. Cell proliferation assessment by EdU incorporation was performed with the Click-iT™ Plus EdU Alexa Fluor™ 647 Flow Cytometry Assay Kit (Thermo Fisher Scientific, C10634) following the manufacturer’s instructions. For detection of MHC-I expression, cells were resuspended as single cells in FACS buffer (PBS containing 2% FBS), stained with anti-human HLA-A,B,C antibody (BioLegend, 311418) for 30 min at room temperature, and washed once with DPBS. All samples were resuspended as single cells in FACS buffer before being acquired by a BD LSRFortessa™ flow cytometer (BD Biosciences).

### Measurement of glucose, ATP, and lactate

Cells were collected and lysed according to the manufacturer’s instructions. Glucose levels were quantified using a glucose assay kit (Nanjing Jiancheng Bioengineering Institute, A154-1-1). Cellular ATP content was measured using an ATP assay kit (Beyotime, S0027), and lactate levels were determined with lactate assay kit (Beyotime, SS0208S). All measurements were performed following the manufacturer’s protocols.

### Non-targeted metabolome analysis

#### Cell harvesting and metabolic quenching

Cells were aspirated to remove culture medium and washed 2 – 3 times with pre-chilled phosphate-buffered saline (PBS) to eliminate residual medium. Cells were trypsinized with 0.25% trypsin-EDTA at 37°C for 2 – 5 min and stopped by complete medium. Cell pellets were collected and immediately quenched with 1 mL of pre-cooled 80% methanol. Samples were resuspended, transferred to cryovials, and stored at −80°C until analysis.

#### Metabolite extraction and sample preparation

Samples were kept on dry ice during processing. For extraction, 100 μL of sample was mixed with 400 μL of extraction solvent (methanol:acetonitrile = 3:1, v/v; pre-cooled to −40°C), vortexed for 5 min, and incubated at 4°C for 2 h. After centrifugation at 12,000 rpm for 15 min at 4°C, supernatants were collected and completely vacuum-dried. Dried extracts were reconstituted in 100 μL of 50% methanol (methanol:water = 1:1, v/v), vortexed for 3 min (4°C, 2000 rpm), and centrifuged at 12,000 rpm for 15 min at 4°C. Supernatants were transferred to LC vials for analysis.

#### UHPLC separation

Chromatographic separation was performed on a Thermo Vanquish ultra-high-performance liquid chromatography (UHPLC) system equipped with an HSS T3 column. The injection volume was 2 μL, column temperature was maintained at 40°C, and the flow rate was 0.3 mL/min. The gradient elution program (mobile phase B) was: 0 – 0.5 min, 2%; 0.5 – 2 min, 2 – 50%; 2 – 5 min, 50 –98%; 5 – 8 min, 98%; 8 – 10 min, 98 – 2%; 10 – 12 min, 2%. Samples were maintained at 8°C in the autosampler. Quality control (QC) samples were interspersed throughout the run to monitor system stability and analytical reproducibility.

#### Mass spectrometry acquisition

Mass spectrometry was conducted using a Thermo Q Exactive HF-X mass spectrometer with electrospray ionization (ESI) operated in both positive (POS) and negative (NEG) ion modes. Source parameters were: sheath gas, 30 L/min; auxiliary gas, 25 L/min; spray voltage, 3.6 kV (POS) and 3.6 kV (NEG); ion transfer RF level, 40; capillary temperature, 325°C; auxiliary gas heater temperature, 300°C. MS/MS data were acquired using stepped normalized collision energies (NCE) of 20, 30, and 40. Data were collected in data-dependent acquisition (DDA) mode (Top N = 10) over an m/z range of 70–1050, with a 12-min acquisition per sample.

#### Data processing, quality assessment, and statistical analysis

Data from POS and NEG ion modes were processed and analyzed separately. QC samples were used to assess instrument performance and batch stability, and PCA was used to evaluate global variation and QC clustering. Unsupervised PCA was conducted using the R package gmodels (v2.18.1). Supervised partial least squares discriminant analysis (PLS-DA) and orthogonal PLS-DA (OPLS-DA) were performed using the R package ropls. Model performance was evaluated by explained variance (R^2^) and predictive ability (Q^2^). Model robustness was assessed by 7-fold cross-validation and 200-iteration permutation testing. Models with Q^2^ > 0.4 were considered acceptable, and Q^2^ > 0.9 were considered indicative of strong predictive performance. Hierarchical cluster analysis (HCA) was performed using the average-linkage method. For visualization, metabolite abundance was z-score-transformed and displayed as heatmaps using the R package pheatmap. Reproducibility among biological replicates was assessed using Pearson correlation coefficients, and correlation heatmaps were generated using pheatmap.

#### Differential metabolite identification and downstream analyses

Differential metabolites were identified based on variable importance in projection (VIP) scores derived from the OPLS-DA model (VIP >= 1) combined with Student’s t test (p < 0.05). Fold changes between groups were calculated and visualized using volcano plots. The top 15 discriminatory metabolites were ranked by VIP scores. Pearson correlation among differential metabolites was calculated using cor and cor.test in R, and correlation matrices were visualized using the R package corrplot. Differential metabolite abundance patterns were visualized by z-score–transformed heatmaps generated with pheatmap. Receiver operating characteristic (ROC) curve analysis was performed using the R package pROC, and the area under the curve (AUC) was calculated to assess discriminatory performance.

#### Pathway annotation and enrichment analysis

Differential metabolites were annotated to the Kyoto Encyclopedia of Genes and Genomes (KEGG) database. Enrichment analysis was performed using a hypergeometric test followed by false discovery rate (FDR) correction, with FDR <= 0.05 considered statistically significant. Metabolite set enrichment analysis (MSEA) was additionally performed using an over-representation analysis (ORA) framework with Fisher’s exact test, based on the Small Molecule Pathway Database (SMPDB) library implemented in the R package MSEAp.

## Figure legends

**Figure S1. Genetic depletion of MTHFD2 enhances tumor susceptibility to CD8+ T cell-mediated killing**.

(A) Validation of MTHFD2 knockdown in A375 cells using two distinct shRNAs (shMTHFD2-1 and shMTHFD2-2) by immunoblotting. β-ACTIN was used as loading control.

(B) Quantification of IFN-γ secretion by T cells in the supernatant (left) and tumor LDH release (right). WT or MTHFD2 knockdown A375 cells were seeded at indicated densities and co-cultured for 24 h with CD8+ T cells expressing 1G4 α95LY TCR. Data are representative of 3 independent experiments and are presented as mean ± SD.

**Figure S2. Drug pretreatment shows limited effects on A498 cells responding to CD8+ T cell-mediated killing**.

Quantification of IFN-γ secretion by T cells in the supernatant (left) and tumor LDH release (right). A498 cells were treated with DMSO, MTHFD2i (5 µM), LTX (2 µM), MTX (2 µM), or 6-MP (100 µM) for 24 h, and then co-cultured for 24 h with CD8+ T cells expressing 1G4 α95LY TCR. Data are representative of 3 independent experiments and are presented as mean ± SD.

**Figure S3. MTHFD2 inhibition has limited effects on HLA and immune checkpoint expression in A375 cells**.

(A) Heatmap of RNA-seq analysis showing transcript levels of HLA and immune checkpoint genes in A375 cells treated with MTHFD2i (5 µM).

(B) Flow cytometric analysis of HLA-A, B, C surface levels on A375 cells treated with MTHFD2i (2 µM or 5 µM) for different durations, with or without IFN-γ (10 ng/mL) supplementation.

**Figure S4. Inhibition of folate-dependent nucleotide synthesis induces cell-cycle arrest in A375 cells, which cannot be rescued by adenine supplementation**.

(A) Cell cycle status of A375 cells treated with MTHFD2i (5 µM) relative to DMSO control, assessed by flow cytometry using the FUCCI2.1 system (AmCyan-hGem/mCherry-hCdt1).

(B) Cell proliferation measured by flow cytometry analysis of EdU incorporation after treatment with DMSO, MTHFD2i (5 µM), LTX (2 µM), MTX (2 µM), or 6-MP (100 µM) with adenine supplementation (5 µM).

(C) IFN-γ secretion in co-culture supernatants. A375 cells seeded at indicated densities were treated with MTHFD2i (5 µM), LTX (2 µM), MTX (2 µM), or 6-MP (100 µM) supplemented with or without adenine (5 µM) for 24 h, and then co-cultured with CD8+ 1G4 α95LY TCR-T cells for another 24 h.

**Figure S5. Inhibition of folate-dependent purine synthesis suppresses mTORC1 signaling, whereas mTOR inhibition does not account for enhanced A375 cell sensitivity to CD8+ T cell killing**.

(A) GSEA of cytoplasmic ribosome-related gene sets in cells treated with MTHFD2i (5 µM) versus DMSO.

(B) Western blot analysis of p-p70-S6K, total p70-S6K, and RHEB protein levels in A375 cells treated with DMSO, MTHFD2i (5 µM), LTX (2 µM), MTX (2 µM), or 6-MP (100 µM). Rescue with adenine (5 µM) or nucleosides (30 μM for cytidine, guanosine, adenosine, and uridine each, and 10 μM for thymidine) was also performed.

(C) Western blot showing the efficiency of mTOR knockdown (mTOR shRNA-1/2) and its effect on p-p70-S6K levels.

(D) IFN-γ secretion levels in co-cultures using mTOR knockdown (shmTOR-1) or control (shGFP) A375 cells at different seeding densities. Cells were co-cultured with CD8+ 1G4 α95LY TCR-T cells for 24 h.

(E) Quantification of IFN-γ secretion. A375 cells were treated with rapamycin for 24 h, and then rapamycin was removed and T cells were added. Co-culture with CD8+ 1G4 α95LY TCR-T cells was performed for another 24h.

(F) Flow cytometric analysis of HLA-A, B, C expression after treatment with indicated concentrations of rapamycin.

(G) Western blot analysis of NY-ESO-1 protein levels after treatment with different concentrations of rapamycin.

**Figure S6. Inhibition of folate-dependent purine synthesis suppresses glycolysis, reduces protein lactylation, and enhances CD8+ T cell-mediated killing in HCT 116 cells**.

(A) Relative LDHA mRNA expression in HCT 116 cells treated with DMSO, MTHFD2i (5 µM), LTX (2 µM), MTX (2 µM), or 6-MP (100 µM) for 24 h.

(B) Protein levels of glycolytic enzymes in HCT 116 cells detected by Immunoblotting. Cells were treated with DMSO, MTHFD2i (5 µM), LTX (2 µM), MTX (2 µM), or 6-MP (100 µM) for 24 h.

(C) Relative intracellular lactate and ATP levels in HCT 116 cells after treatment with DMSO, MTHFD2i (5 µM), LTX (2 µM), MTX (2 µM), or 6-MP (100 µM) for 24 h.

(D) Relative intracellular glucose levels in HCT 116 cells after treatment with DMSO, MTHFD2i (5 µM), LTX (2 µM), MTX (2 µM), or 6-MP (100 µM) for 24 h.

(E) Relative intracellular lactate and ATP levels in WT and MTHFD2-depleted A375 cells.

(F) IFN-γ secretion in co-culture supernatants. HCT 116 cells seeded at indicated densities were treated with DMSO, MTHFD2i (5 µM), 2-DG (50 µM), or Lonidamine (50 µM) for 24 h, then co-cultured with CD8+ MAGE-A3 94a-14 TCR-T cells for another 24 h.

(G) IFN-γ secretion (upper panels) and tumor LDH release (lower panels) in co-culture assays. HCT 116 cells seeded at indicated densities were treated with DMSO, MTHFD2i (5 µM), LTX (2 µM), MTX (2 µM), or 6-MP (100 µM) for 24 h, followed by co-culture with MAGE-A3 94a-14 TCR-T cells in the absence or presence of exogenous lactate (10 mM).

(H) CCK-8 assay showing relative viability of A375 cells treated with MTHFD2i (5 µM), LTX (2 µM), MTX (2 µM) for 24 h. Then cells were untouched (No wash), washed with DPBS and replaced with fresh medium (Wash), or washed and replaced with fresh medium containing lactate (10 mM) (Wash + lactate).

(I) Immunoblotting of global protein lactylation in HCT 116 cells treated with DMSO, MTHFD2i (5 µM), LTX (2 µM), MTX (2 µM), or 6-MP (100 µM) for 24 h in the absence (Drug) or presence (Drug + lactate) of sodium L-lactate (10 mM) and detected for global protein lactylation.

**Figure S7. Relative AICAR abundance in metabolomics analysis of drug-treated A375 cells**.

AICAR abundance in metabolomics data. Relative AICAR levels measured by LC-MS in A375 cells treated with MTHFD2i (5 µM, left) or LTX (2 µM, right) compared with DMSO control. Each dot represents an independent biological replicate.

## Notes

https://www.ebi.ac.uk/metabolights/MTBLS13434

### Competing Interest Statement

The authors have declared no competing interest.

## References

1 Curtis, L. T., Sebens, S. & Frieboes, H. B. Modeling of tumor response to macrophage and T lymphocyte interactions in the liver metastatic microenvironment. Cancer Immunol Immunother 70, 1475–1488, doi:10.1007/s00262-020-02785-4 (2021).

2 Altea-Manzano, P., Decker-Farrell, A., Janowitz, T. & Erez, A. Metabolic interplays between the tumour and the host shape the tumour macroenvironment. Nat Rev Cancer 25, 274–292, doi:10.1038/s41568-024-00786-4 (2025).

3 Korsmo, H. W. & Jiang, X. One carbon metabolism and early development: a diet-dependent destiny. Trends Endocrinol Metab 32, 579–593, doi:10.1016/j.tem.2021.05.011 (2021).

4 Ducker, G. S. & Rabinowitz, J. D. One-Carbon Metabolism in Health and Disease. Cell Metab 25, 27–42, doi:10.1016/j.cmet.2016.08.009 (2017).

5 Yang, M. & Vousden, K. H. Serine and one-carbon metabolism in cancer. Nat Rev Cancer 16, 650–662, doi:10.1038/nrc.2016.81 (2016).

6 Aury-Landas, J. et al. Anti-inflammatory and chondroprotective effects of the S-adenosylhomocysteine hydrolase inhibitor 3-Deazaneplanocin A, in human articular chondrocytes. Sci Rep 7, 6483, doi:10.1038/s41598-017-06913-6 (2017).

7 Nouzova, M., Michalkova, V., Ramirez, C. E., Fernandez-Lima, F. & Noriega, F. G. Inhibition of juvenile hormone synthesis in mosquitoes by the methylation inhibitor 3-deazaneplanocin A (DZNep). Insect Biochem Mol Biol 113, 103183, doi:10.1016/j.ibmb.2019.103183 (2019).

8 Bose, S. & Le, A. Glucose Metabolism in Cancer. Adv Exp Med Biol 1063, 3–12, doi:10.1007/978-3-319-77736-8_1 (2018).

9 Amelio, I., Cutruzzola, F., Antonov, A., Agostini, M. & Melino, G. Serine and glycine metabolism in cancer. Trends Biochem Sci 39, 191–198, doi:10.1016/j.tibs.2014.02.004 (2014).

10 Chang, C. H. et al. Metabolic Competition in the Tumor Microenvironment Is a Driver of Cancer Progression. Cell 162, 1229–1241, doi:10.1016/j.cell.2015.08.016 (2015).

11 TeSlaa, T., Ralser, M., Fan, J. & Rabinowitz, J. D. The pentose phosphate pathway in health and disease. Nat Metab 5, 1275–1289, doi:10.1038/s42255-023-00863-2 (2023).

12 Lane, A. N. & Fan, T. W. Regulation of mammalian nucleotide metabolism and biosynthesis. Nucleic Acids Res 43, 2466–2485, doi:10.1093/nar/gkv047 (2015).

13 Skinner, O. S. et al. Salvage of ribose from uridine or RNA supports glycolysis in nutrient-limited conditions. Nat Metab 5, 765–776, doi:10.1038/s42255-023-00774-2 (2023).

14 Nwosu, Z. C. et al. Uridine-derived ribose fuels glucose-restricted pancreatic cancer. Nature 618, 151–158, doi:10.1038/s41586-023-06073-w (2023).

15 Liberti, M. V. & Locasale, J. W. The Warburg Effect: How Does it Benefit Cancer Cells? Trends Biochem Sci 41, 211–218, doi:10.1016/j.tibs.2015.12.001 (2016).

16 Frey, A. B. & Monu, N. Cancer-Induced Signaling Defects in Antitumor T Cells. Immunol Rev. 222, 192–205, doi:10.1111/j.1600-065X.2008.00606.x (2008).

17 Egen, J. G., Ouyang, W. & Wu, L. C. Human Anti-tumor Immunity: Insights from Immunotherapy Clinical Trials. Immunity 52, 36–54, doi:10.1016/j.immuni.2019.12.010 (2020).

18 Hou, M., Ji, L., Li, D., Xiao, Q. & Hu, X. Serum starvation-induced cholesterol reduction increases melanoma cell susceptibility to cytotoxic T lymphocyte killing. Sci Rep 15, 18364, doi:10.1038/s41598-025-00586-2 (2025).

19 Fang, D. D. et al. MDM2 inhibitor APG-115 synergizes with PD-1 blockade through enhancing antitumor immunity in the tumor microenvironment. J Immunother Cancer 7, 327, doi:10.1186/s40425-019-0750-6 (2019).

20 Zhou, X. et al. Pharmacologic Activation of p53 Triggers Viral Mimicry Response Thereby Abolishing Tumor Immune Evasion and Promoting Antitumor Immunity. Cancer Discov 11, 3090–3105, doi:10.1158/2159-8290.CD-20-1741 (2021).

21 Dekhne, A. S., Hou, Z., Gangjee, A. & Matherly, L. H. Therapeutic Targeting of Mitochondrial One-Carbon Metabolism in Cancer. Mol Cancer Ther 19, 2245–2255, doi:10.1158/1535-7163.MCT-20-0423 (2020).

22 Zhao, X. et al. Tuning T cell receptor sensitivity through catch bond engineering. Science 376, eabl5282, doi:10.1126/science.abl5282 (2022).

23 Clare, C. E., Brassington, A. H., Kwong, W. Y. & Sinclair, K. D. One-Carbon Metabolism: Linking Nutritional Biochemistry to Epigenetic Programming of Long-Term Development. Annu Rev Anim Biosci 7, 263–287, doi:10.1146/annurev-animal-020518-115206 (2019).

24 Chandel, N. S. Nucleotide Metabolism. Cold Spring Harb Perspect Biol 13, doi:10.1101/cshperspect.a040592 (2021).

25 Karran, P. & Attard, N. Thiopurines in current medical practice: molecular mechanisms and contributions to therapy-related cancer. Nat Rev Cancer 8, 24–36, doi:10.1038/nrc2292 (2008).

26 Cornish, J. S., Wirthgen, E. & Dabritz, J. Biomarkers Predictive of Response to Thiopurine Therapy in Inflammatory Bowel Disease. Front Med (Lausanne) 7, 8, doi:10.3389/fmed.2020.00008 (2020).

27 Karim, H., Ghalali, A., Lafolie, P., Vitols, S. & Fotoohi, A. K. Differential role of thiopurine methyltransferase in the cytotoxic effects of 6-mercaptopurine and 6-thioguanine on human leukemia cells. Biochem Biophys Res Commun 437, 280–286, doi:10.1016/j.bbrc.2013.06.067 (2013).

28 Tran, D. H. et al. De novo and salvage purine synthesis pathways across tissues and tumors. Cell 187, 3602–3618 e3620, doi:10.1016/j.cell.2024.05.011 (2024).

29 Wilde, B. R. et al. FH Variant Pathogenicity Promotes Purine Salvage Pathway Dependence in Kidney Cancer. Cancer Discov 13, 2072–2089, doi:10.1158/2159-8290.CD-22-0874 (2023).

30 Nishimura, K. et al. APC(CDH1) targets MgcRacGAP for destruction in the late M phase. PLoS One 8, e63001, doi:10.1371/journal.pone.0063001 (2013).

31 Emmanuel, N. et al. Purine Nucleotide Availability Regulates mTORC1 Activity through the Rheb GTPase. Cell Rep 19, 2665–2680, doi:10.1016/j.celrep.2017.05.043 (2017).

32 Hoxhaj, G. et al. The mTORC1 Signaling Network Senses Changes in Cellular Purine Nucleotide Levels. Cell Rep 21, 1331–1346, doi:10.1016/j.celrep.2017.10.029 (2017).

33 Saxton, R. A. & Sabatini, D. M. mTOR Signaling in Growth, Metabolism, and Disease. Cell 168, 960–976, doi:10.1016/j.cell.2017.02.004 (2017).

34 Saucedo, L. J. et al. Rheb promotes cell growth as a component of the insulin/TOR signalling network. Nat Cell Biol 5, 566–571, doi:10.1038/ncb996 (2003).

35 Stocker, H. et al. Rheb is an essential regulator of S6K in controlling cell growth in Drosophila. Nat Cell Biol 5, 559–565, doi:10.1038/ncb995 (2003).

36 Brand, A. et al. LDHA-Associated Lactic Acid Production Blunts Tumor Immunosurveillance by T and NK Cells. Cell Metab 24, 657–671, doi:10.1016/j.cmet.2016.08.011 (2016).

37 Angelin, A. et al. Foxp3 Reprograms T Cell Metabolism to Function in Low-Glucose, High-Lactate Environments. Cell Metab 25, 1282–1293 e1287, doi:10.1016/j.cmet.2016.12.018 (2017).

38 Dietl, K. et al. Lactic acid and acidification inhibit TNF secretion and glycolysis of human monocytes. J Immunol 184, 1200–1209, doi:10.4049/jimmunol.0902584 (2010).

39 Zong, Z., Ren, J., Yang, B., Zhang, L. & Zhou, F. Emerging roles of lysine lactyltransferases and lactylation. Nat Cell Biol 27, 563–574, doi:10.1038/s41556-025-01635-8 (2025).

40 Huang, Z. W. et al. STAT5 promotes PD-L1 expression by facilitating histone lactylation to drive immunosuppression in acute myeloid leukemia. Signal Transduct Target Ther 8, 391, doi:10.1038/s41392-023-01605-2 (2023).

41 Tong, H. et al. Dual impacts of serine/glycine-free diet in enhancing antitumor immunity and promoting evasion via PD-L1 lactylation. Cell Metab 36, 2493–2510 e2499, doi:10.1016/j.cmet.2024.10.019 (2024).

42 Schjoldager, K. T., Narimatsu, Y., Joshi, H. J. & Clausen, H. Global view of human protein glycosylation pathways and functions. Nat Rev Mol Cell Biol 21, 729–749, doi:10.1038/s41580-020-00294-x (2020).

43 Shang, M. et al. The folate cycle enzyme MTHFD2 induces cancer immune evasion through PD-L1 up-regulation. Nat Commun 12, 1940, doi:10.1038/s41467-021-22173-5 (2021).

44 Strefeler, A., Blanco-Fernandez, J. & Jourdain, A. A. Nucleosides are overlooked fuels in central carbon metabolism. Trends Endocrinol Metab 35, 290–299, doi:10.1016/j.tem.2024.01.013 (2024).

45 Choi, J. W. et al. Uridine protects cortical neurons from glucose deprivation-induced death: possible role of uridine phosphorylase. J Neurotrauma 25, 695–707, doi:10.1089/neu.2007.0409 (2008).

46 Song, C., Li, Q., Zhang, J. & Hu, W. Uridine Phosphorylase 1 as a Biomarker Associated with Glycolysis in Acute Lung Injury. Inflammation, doi:10.1007/s10753-025-02270-z (2025).

47 Qiao, J. et al. The Pentose Phosphate Pathway: From Mechanisms to Implications for Gastrointestinal Cancers. Int J Mol Sci 26, doi:10.3390/ijms26020610 (2025).

48 Tozzi, M. G., Camici, M., Mascia, L., Sgarrella, F. & Ipata, P. L. Pentose phosphates in nucleoside interconversion and catabolism. FEBS J 273, 1089–1101, doi:10.1111/j.1742-4658.2006.05155.x (2006).

49 Hosomi, S., Tara, H., Terada, T. & Mizoguchi, T. Inhibitory effect of 5-phosphoribosyl 1-pyrophosphate and ADP on the nonoxidative pentose phosphate pathway activity. Biochem Med Metab Biol 42, 52–59, doi:10.1016/0885-4505(89)90040-6 (1989).

50 Antoniewicz, M. R. A guide to (13)C metabolic flux analysis for the cancer biologist. Exp Mol Med 50, 1–13, doi:10.1038/s12276-018-0060-y (2018).

51 Metallo, C. M., Walther, J. L. & Stephanopoulos, G. Evaluation of 13C isotopic tracers for metabolic flux analysis in mammalian cells. J Biotechnol 144, 167–174, doi:10.1016/j.jbiotec.2009.07.010 (2009).

52 Sugiura, A. et al. MTHFD2 is a metabolic checkpoint controlling effector and regulatory T cell fate and function. Immunity 55, 65–81 e69, doi:10.1016/j.immuni.2021.10.011 (2022).

