## Supplementary figures and images for "Targeting folate-dependent purine synthesis sensitizes melanoma cells to immune attack through suppressing glycolysis"

### FigureS1

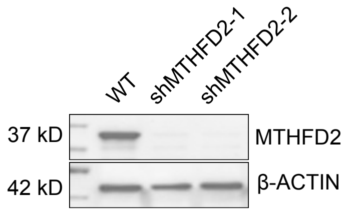

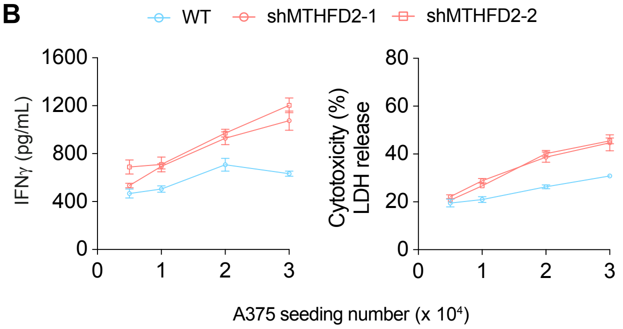

### FigureS2

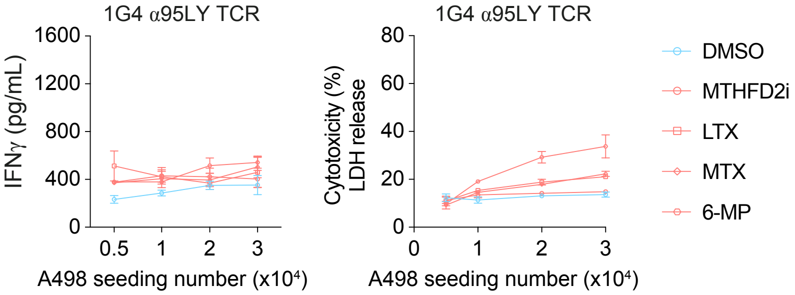

### FigureS3

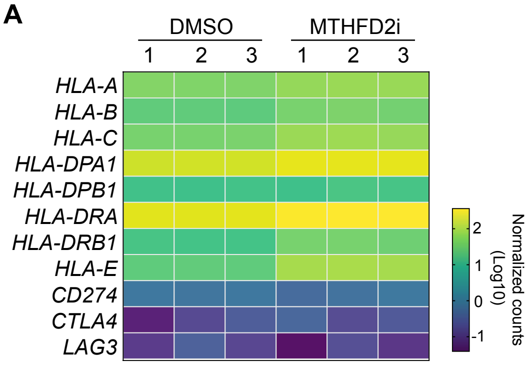

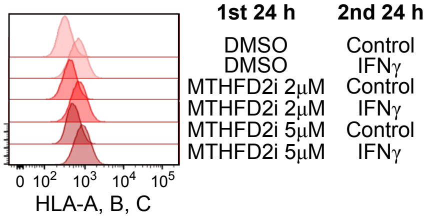

### FigureS4

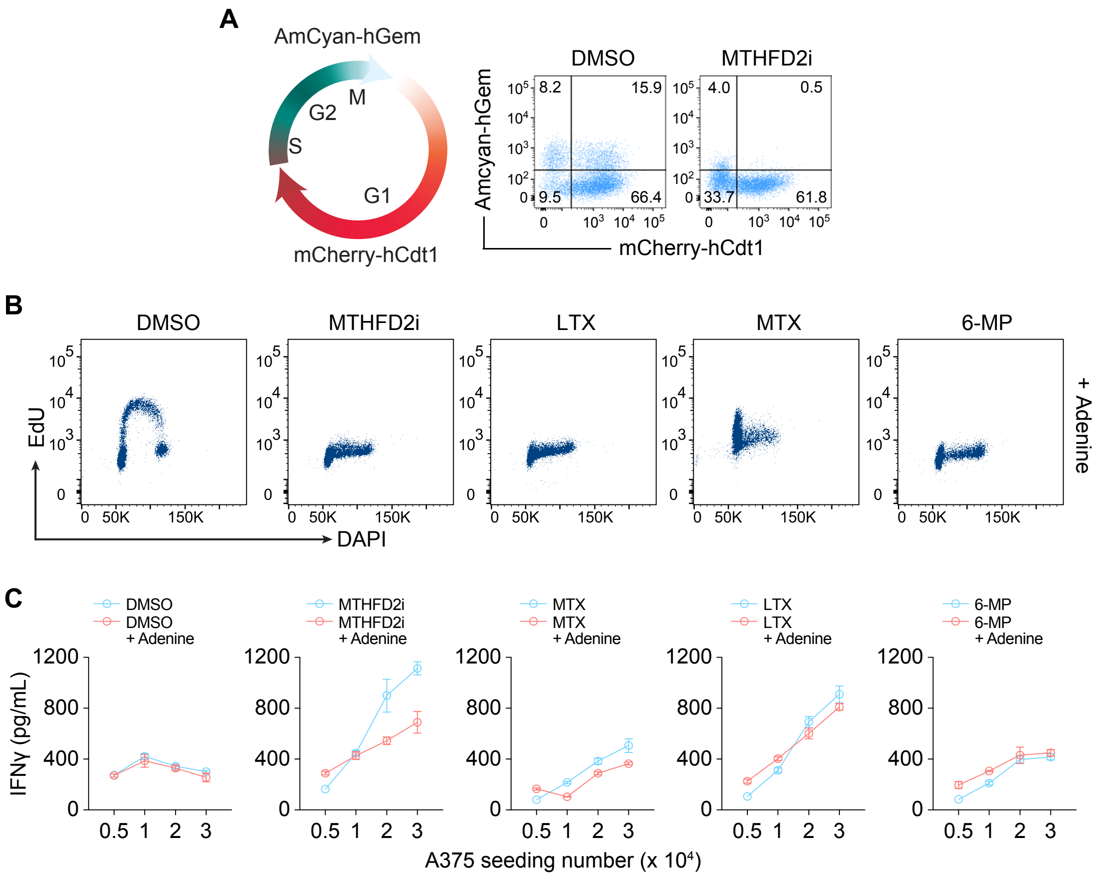

### FigureS5

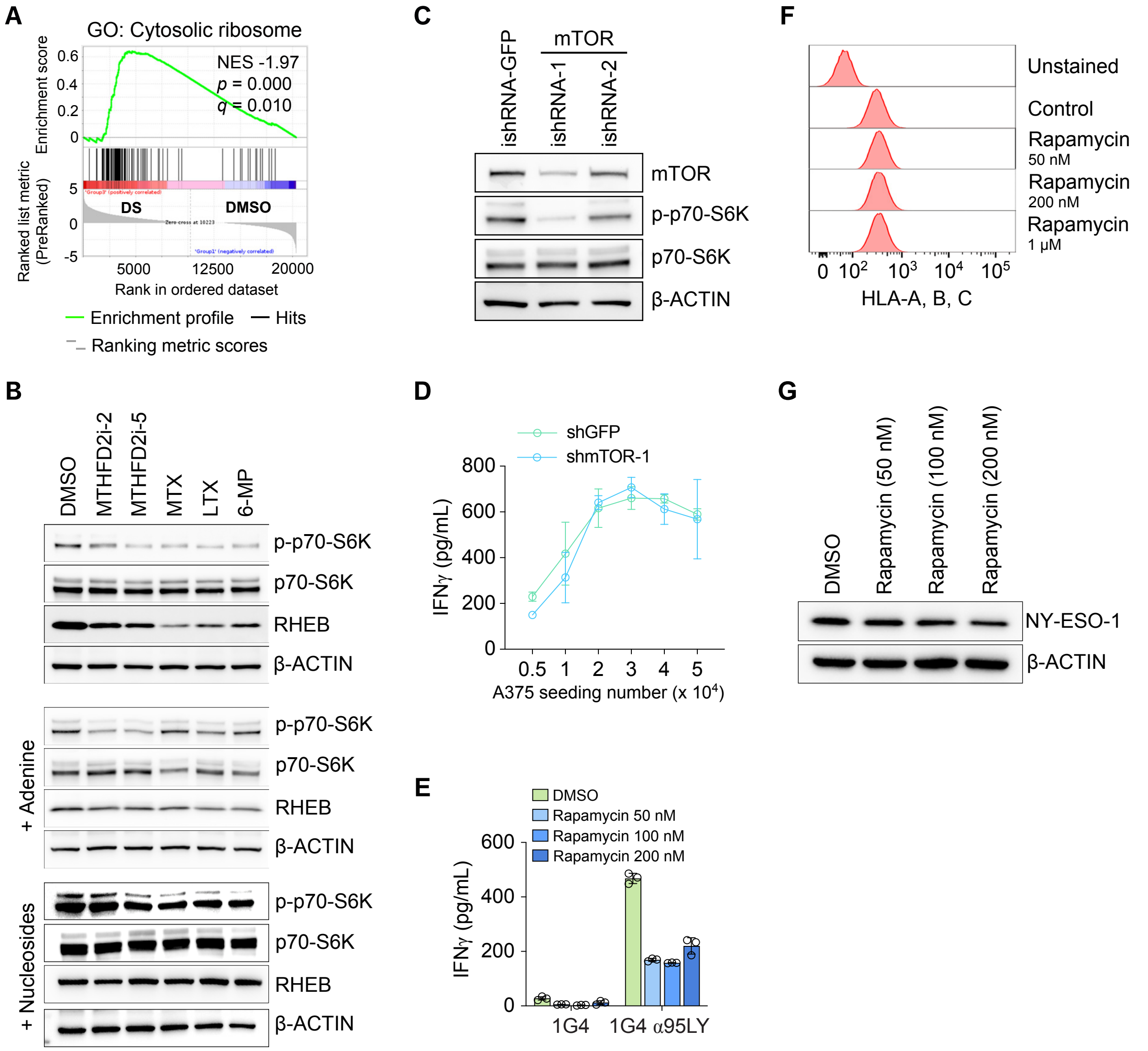

### FigureS6

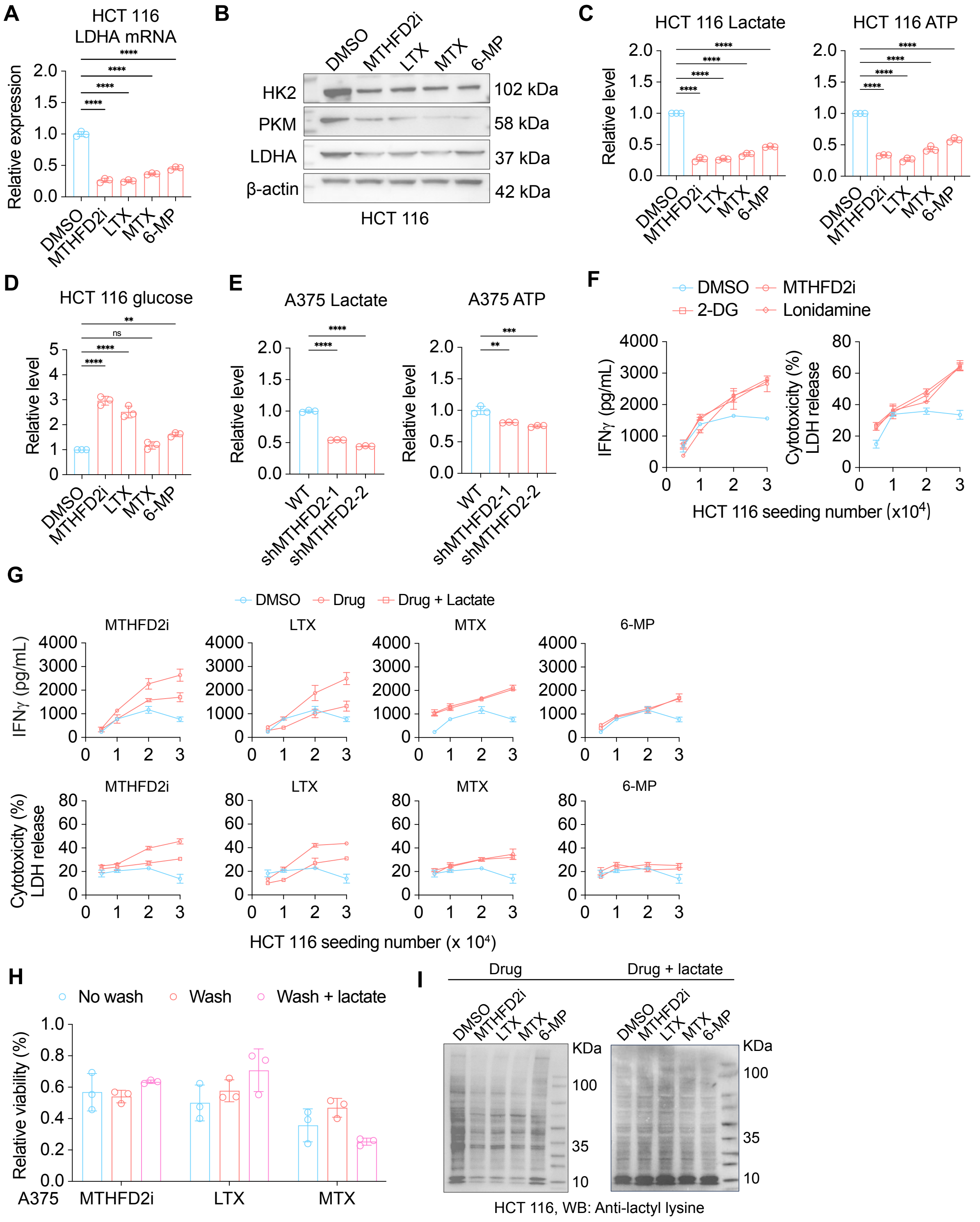

### FigureS7

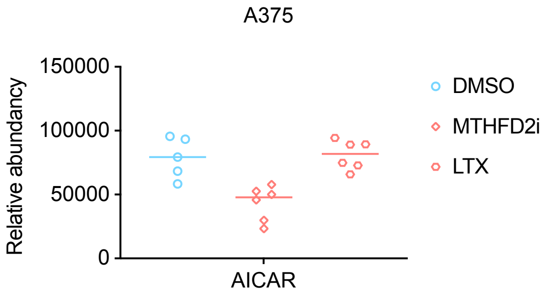
